# *In silico* analysis of *cis*-elements and identification of transcription factors putatively involved in the regulation of the OAS cluster genes *SDI1* and *SDI2*

**DOI:** 10.1101/2021.03.30.437644

**Authors:** Apidet Rakpenthai, Anastasia Apodiakou, Sarah J. Whitcomb, Rainer Hoefgen

## Abstract

*A. thaliana sulfur deficiency-induced 1* and *sulfur deficiency-induced 2* (*SDI1* and *SDI2*) are involved in partitioning sulfur among metabolite pools during sulfur deficiency and their transcription is strongly induced by this condition. However, little is currently known about the *cis*- and *trans*-factors that regulate *SDI* expression. To identify potential transcription factors and DNA sequence element regulators of *SDI* expression we performed a comparative *in silico* analysis of their promoter sequences cataloguing known and potentially new *cis*-elements. We further screened an arrayed library of Arabidopsis transcription factors (TF) for binding to the *SDI1* and *SDI2* promoters. In total 14 candidate TF regulators of *SDI*s were identified with yeast-one-hybrid analyses, of which five bound to both promoters, 4 were specific to *SDI1,* and 5 were specific *SDI2*. Direct association between particular *cis*-elements in these promoter regions and specific TFs was established via electrophoretic mobility shift assays. SLIM1 was shown to bind SURE *cis*-element(s) in the proximal promoter region of both *SDI1* and *SDI2*. The bZIP core *cis*-element in the proximal promoter region of *SDI2* was shown to be important for bZIP16, bZIP44, and HYH binding. GBF1 was shown to bind the E-box in the proximal promoter region of *SDI2*. Additionally, we performed a meta-analysis of expression changes of these 14 TF candidates in a variety of conditions that alter *SDI* expression. These data will allow for more detailed future analysis of the molecular factors required for transcriptional regulation of *SDI*s under a range of physiological and metabolic conditions, apart from sulfur deficiency.

## INTRODUCTION

Sulfur (S) is an essential macronutrient for plant growth and development. S is predominantly taken up from the soil in the form of sulfate and then enzymatically reduced and assimilated into S-containing metabolites including cysteine, methionine, *S*-adenosylmethionine (SAM), glutathione, cofactors such as coenzymeA, glucosinolates (GSL), and sulfolipids. These compounds play pivotal roles for essential biological processes in plants, namely redox control and detoxification by glutathione, methylation and ethylene biosynthesis by SAM, and anti-herbivory defense by GSL, among others. S-limited growth conditions elicit a network of responses, including upregulation of sulfate transporters located in the plasma and tonoplast membranes, increased activity of enzymatic steps involved in sulfate reduction and assimilation, and changes in S-partitioning between major metabolite pools (Saito et al., 2000, 2004; Hoefgen and Nikiforova, 2008; Lewandowska and Sirko, 2008; Takahashi et al., 2011; Chan et al., 2019; Kopriva et al., 2019).

*Sulfur deficiency-induced 1* (*SDI1*) is among the genes most strongly induced by S-limitation (Maruyama-Nakashita et al., 2006; Dietzen et al., 2020). Several studies using knockout and overexpression lines have shown *SDI1* to be important for partitioning of sulfur among S-metabolite pools, especially under low-S. In *Brassicale* species such as Arabidopsis, GSL account for a large fraction of the total S-containing metabolites (Falk et al., 2007), and when S becomes limiting, plants repress GSL production (Zhao et al., 1994; Aghajanzadeh et al., 2014), in part through the activity of SDI1 (Aarabi et al., 2016, 2020). An important role for SDI1 in regulating sulfate assimilation is suggested by the higher internal sulfate content in *sdi1* knockout plants compared to wild type during S-stress (Howarth et al., 2009; Aarabi et al., 2016), perhaps due to reduced efflux of vacuolar sulfate and/or reduced assimilation in the absence of SDI1. *Sulfur deficiency-induced 2* (*SDI2*), a homolog of *SDI1* with 60% protein sequence identity, similar protein length and position of its TPR domain, is also upregulated by low-S (Maruyama-Nakashita et al., 2006; Aarabi et al., 2016; Dietzen et al., 2020). In Arabidopsis, *SDI1* and *SDI2* have similar effects on GSL accumulation in seedlings under full-nutrient (FN) and –S conditions. Further, phenotypes in *sdi1sdi2* double knockout lines tend to be stronger than in the single knockout parents, also suggesting significant functional redundancy of SDI1 and SDI2, at least in these tissues and conditions (Aarabi et al., 2016).

Multiple studies have characterized transcriptional responses to S-deficiency (Ohkama et al., 2002; Maruyama-Nakashita et al., 2003; Nikiforova et al., 2003; Goda et al., 2008; Iyer-Pascuzzi et al., 2011; Forieri et al., 2017). However, our understanding of how those responses, including *SDI* induction, are regulated on the molecular level is still very patchy. Mutations in *Sulfur limitation 1* (SLIM1) affect expression of most low S responsive genes in Arabidopsis roots (Kawashima et al., 2011; Maruyama-Nakashita et al., 2006; Dietzen et al., 2020). However, the promoters of only approximately 15% of those genes contained a SLIM1 binding peak by DAP-seq (O’Malley et al., 2016). The transcript level of several TFs involved in auxin signaling (IAA13, IAA28, and ARF-2) have been shown to respond strongly to sulfur starvation, and mutations in these genes result in altered S-compound contents (Falkenberg et al., 2008; Hoefgen et al., 2017), but no direct targets of these TFs have thus far been identified in the context of S-limitation responses. Of the eight TFs known to have at least one direct target in the S-deficiency transcriptional response (HY5, MYB28, MYB29, MYB34, MYB51, MYB76, MYB122, and SLIM1), none are strongly affected on the transcript level by insufficient S (Maruyama-Nakashita et al., 2006; Yatusevich et al., 2010; Lee et al., 2011).

Three *cis*-regulatory elements involved in sulfur response have been identified so far. The sulfur-responsive element (SURE) was identified in the promoter of *sulfate transporter 1;1*. The core sequence of the SURE element (GAGAC) is sufficient to drive sulfur-deprivation dependent transcription of a reporter gene and is found in the promoters of many sulfur-responsive genes (Maruyama-Nakashita et al., 2005), but it is currently unknown which transcriptional regulators bind this sequence. The UPE box (AG[G/A]T[T/A]CATTGAA[T/C]CT[A/G]GAC[A/G]) contains two analogous TEIL binding site (TEBS) sequences (A[T/C]G[A/T]A[C/T]CT) and is present in the promoters of several sulfur deficiency-responsive genes. Mutation of the UPE box affects the yeast-one-hybrid (Y1H) binding strength of SLIM1 to the promoter of a direct target, *LSU1* (Wawrzyńska et al., 2010). SURE, UPE, and TEBS *cis*-elements are frequently located in the proximalń (*prox*) promoter region of S-responsive genes, including *SDI1* and *SDI2* (Wawrzyńska et al., 2010; Aarabi et al., 2015), a promoter region that tends to be enriched in TF binding sites (TFBS) (Yu et al., 2016). Furthermore, analysis of *in vitro* genome-wide TF binding sites by DNA affinity purification sequencing (DAP-seq) identified a putative SLIM1-binding motif ([T/A]G[T/A]A[T/C]C[T/A][A/G]GAC[A/G]) that shares significant sequence similarity with the UPE box (O’Malley et al., 2016).

To our knowledge, SDI function has only been studied in the context of S-metabolism. However, their moderate expression level at many developmental stages and in many conditions (Figure S1) suggests that these genes are likely to have other functions in addition to responses to S-deficiency. In fact, *SDI* expression can also be regulated by oxidative stress (Scarpeci et al., 2008 ArrayExpress: E-ATMX-28; Sharma et al., 2013 GEO: GSE41963; Marmiroli et al., 2014 GEO: GSE53989), high light-stress (Shao et al., 2013 GEO: GSE49596; Van Aken et al., 2013 GEO: GSE46107; Glasser et al., 2014 ArrayExpress: E-MTAB-1344), phytohormones such as jasmonic acid, abscisic acid, and auxin (Pauwels et al., 2008 ArrayExpress: E-ATMX-13; Umezawa et al., 2013 ArrayExpress: E-MEXP-3713; Delker et al., 2010 GEO: GSE18975), sudden light shifts (Caldana et al., 2011 ArrayExpress: E-MTAB-375; Hubberten et al., 2012), circadian rhythms (Espinoza et al., 2010 ArrayExpress: E-MEXP-2526; Hubberten et al., 2012; Rugnone et al., 2013 GEO: GSE46621), and by hypoxia (van Dongen et al., 2009 GEO: GSE11558; Yang et al., 2011 GEO: GSE27475). Interestingly shifting plants from light to dark results in a strong but transient accumulation of O-acetylserine (OAS) (Caldana et al., 2011), a compound that accumulates during S-starvation (Kimura et al., 2006; Maruyama-Nakashita et al., 2004), is a direct precursor of cysteine, and regulates accumulation of sulfur containing seed storage-protein genes (Kim et al., 1999). In fact, exogenous OAS treatment (Hirai et al., 2003) and endogenous induction of serine◻O◻acetyltransferase, the enzyme that synthesizes OAS (Hubberten et al., 2012), also strongly induce *SDI*s even in sufficient-S conditions.

Although expression of both *SDI1* and *SDI2* are influenced by the abovementioned conditions, there are significant differences in their expression patterns. For instance, expression of *SDI1* tends to be substantially lower than *SDI2* in all developmental stages/tissues of Arabidopsis except in mature siliques and senescent leaves (Zimmermann et al., 2005) (Figure S1). In contrast, *SDI1* tends to be more strongly induced than *SDI2* by the conditions described above, including S-deficiency. Further indicating differences in transcriptional regulation of *SDI1* and *SDI2*, expression of *SDI2* was found to be similar in *slim1* and WT under both FN and S-deficient conditions, while *SDI1* was strongly mis-regulated in *slim1* mutants (Maruyama-Nakashita et al., 2006 GEO: GSE4455; Dietzen et al., 2020 GEO: GSE157765).

Due to the demonstrated importance of SDIs in sulfur-deficiency responses and hints of involvement in other physiological contexts, here we investigate *SDI* promoter sequence elements and identify candidate TF regulators of *SDI1* and *SDI2* expression. Our Y1H screening identified 14 TFs that can bind to *SDI* promoter regions, and for the four strongest binders we utilized an electrophoretic mobility shift assay (EMSA) to support TF binding to particular short sequence elements in the *SDI* promoters. Notably, our Y1H screening with the *SDI1* and *SDI2* promoters resulted in only partially overlapping set of TFs, which may reflect some of the molecular mechanisms that result in differential expression of these homologs.

## RESULTS

### *In silico* analysis of *SDI* promoter sequences

As a first step towards understanding transcriptional regulation of the *SDI* genes, we used the Arabidopsis *cis*-regulatory element database (AtcisDB) to search the promoters of *SDI1* and *SDI2* for TFBS and their corresponding TFs/TF families. AtcisDB did not contain any experimentally validated TF-TFBS pairs in these promoter regions (2000-bp upstream of the translation start site), but it did contain predicted binding sites for a variety of TF families, such as ABI3VP1, bHLH, bZIP, EIL, HB, LFY, MYB, and WRKY (Figure 1A and Table 1). Notably, specific TFs from the HB, MYB, and WRKY families are regulated by sulfur deficiency in Arabidopsis: HAT14 (HB family), MYB31, MYB45, MYB52, MYB53, MYB54, MYB71, MYB75, MYB93, and WRKY56 are upregulated while MYB29 is downregulated by –S (Bielecka et al., 2015). Furthermore, because of the presence of a RY *cis*-element in the *SDI1* promoter (Figure 1A and Table 1) and the significance of links between OAS and sulfur regulation of seed storage-protein composition (Kim et al., 1999), major TFs from the B3 family, namely FUS3, ABI3, LEC1 and LEC2, which are involved in the regulation of seed storage protein gene expression in Arabidopsis (Luerssen et al., 1998; Stone et al., 2001), may be relevant to *SDI* expression.

**Figure 1:**
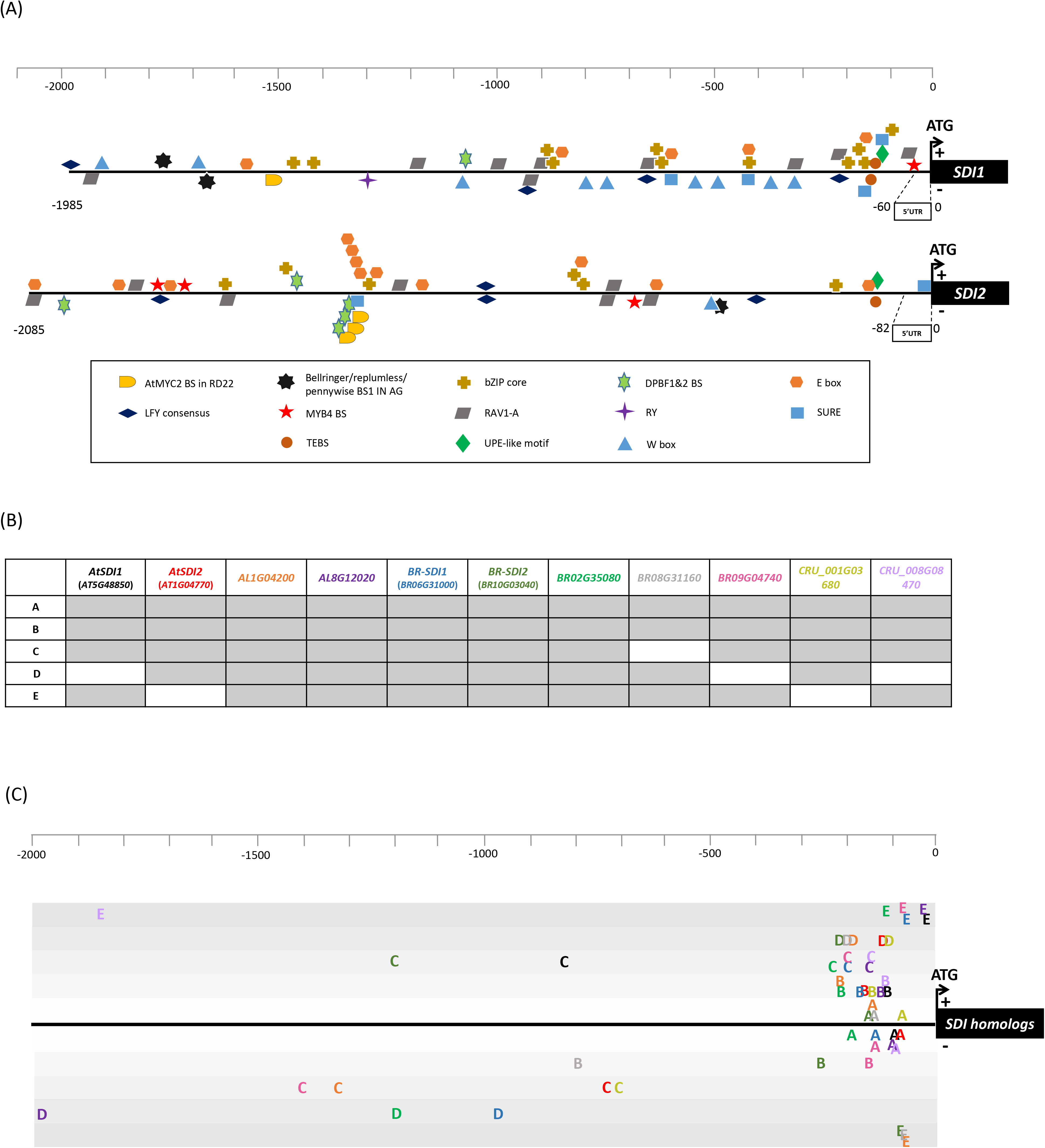
Schematic diagram of cis-elements and putative sequence motifs in *SDI1* and *SDI2* promoters. (**A**) The positions of cis-elements from AtcisDB and PLACE databases as well as cis-elements relevant for sulfur starvation responses are shown in the promoter region of *SDI1* (including the 60-bp 5′ UTR) and *SDI2* (including the 82-bp 5′ UTR). (**B**, **C**) Five putative motifs (represented by single letter codes A through E) were identified by MEME suite 5.1.1 based on promoter sequence conservation among SDI-homolog genes in four Brassicaceae species, At: *Arabidopsis thaliana*, AL: *Arabidopsis lyrata*, BR: *Brassica rapa*, and CRU: *Capsella rubella*). In **B,** the presence (gray fill) or absence (white fill) of each motif in the promoters of the 11 *SDI*-homolog genes is indicated. Each motif occurrence in these promoters is shown in **C**. All motifs found in a given promoter are assigned the same color as the gene name in **B**. (**A**, **C**) *Cis*-elements and putative motifs found on the (+) sense strand are shown above the promoter line while those found on the (−) anti-sense strand are shown below.

**Table 1:**
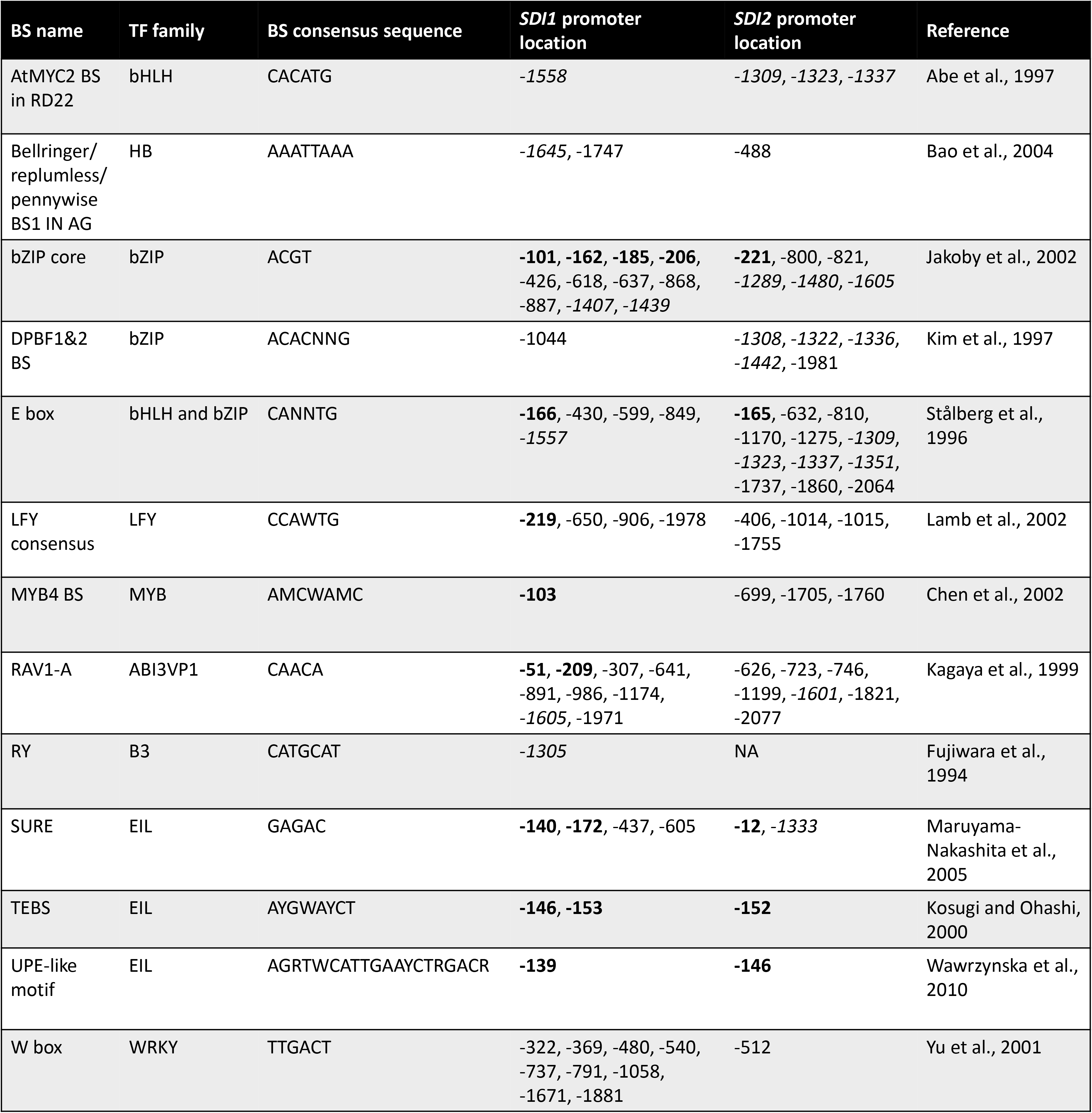
Cis-elements identified by AtcisDB and PLACE databases in *SDI1* and *SDI2* promoters. The cis-elements identified by AtcisDB and PLACE are presented here in binding site (BS) - TF family pairs. Ambiguous bases in the BS sequences are notated with their IUPAC code: M for either A or C; N for any base; R for either A or G; W for either A or T; Y for either C or T. The precise locations of the cis-elements are relative to the ATG start codon of *SDI1* and *SDI2*. Bold- and italic-fonts indicate positions within the regions we define as *prox*- and *dist*-of the *SDI* promoters, respectively.

In addition to those TFBS mapped with AtcisDB, the *SDI1* and *SDI2* promoters also contain several well established *cis*-elements that may be relevant for transcriptional regulation of these genes. Specifically, SURE, which contributes to transcriptional regulation of some sulfur-responsive genes (Maruyama-Nakashita et al., 2005), is present in four and two copies in the promoters of *SDI1* and *SDI2*, respectively. Further, the proximal promoter region of both *SDI1* and *SDI2* contain TEBS sequences and a motif very similar to the UPE box, a SLIM1 binding site (Kosugi and Ohashi, 2000; Wawrzyńska et al., 2010) (Figure 1A and Table 1), which we call the UPE-like motif.

We also searched these promoters for novel sequence motifs. Phylogenetic analysis performed by Aarabi and colleagues (2016) identified nine putative homologs of *AtSDI1* and *AtSDI2* in three other Brassicaceae species (*Arabidopsis lyrata*, *Brassica rapa*, and *Capsella rubella*). The promoter sequences (2000-bp upstream of the translation start site) of these 11 genes were used as input for novel motif identification using MEME suite 5.1.1. Five motifs 10-12 nucleotides in length and with an E-value ≤ 0.05 were identified in the promoters and named A through E (Figure 1B). Their consensus sequences and locations in *SDI*-homolog promoters are shown in Table 2 and Figure 1C. Motifs A, B, C, and E are present in the promoter of *AtSDI1*, whereas the *AtSDI2* promoter contains motifs A, B, C, and D. Interestingly, the promoters of all 11 Brassicaceae *SDI*-homologs contain motif A within 250-bp of the translation start site (Figure 1C). Such conservation suggests that motif A may function as a *cis*-element controlling transcriptional regulation of SDI family genes in *Brassicaceae*.

**Table 2:**
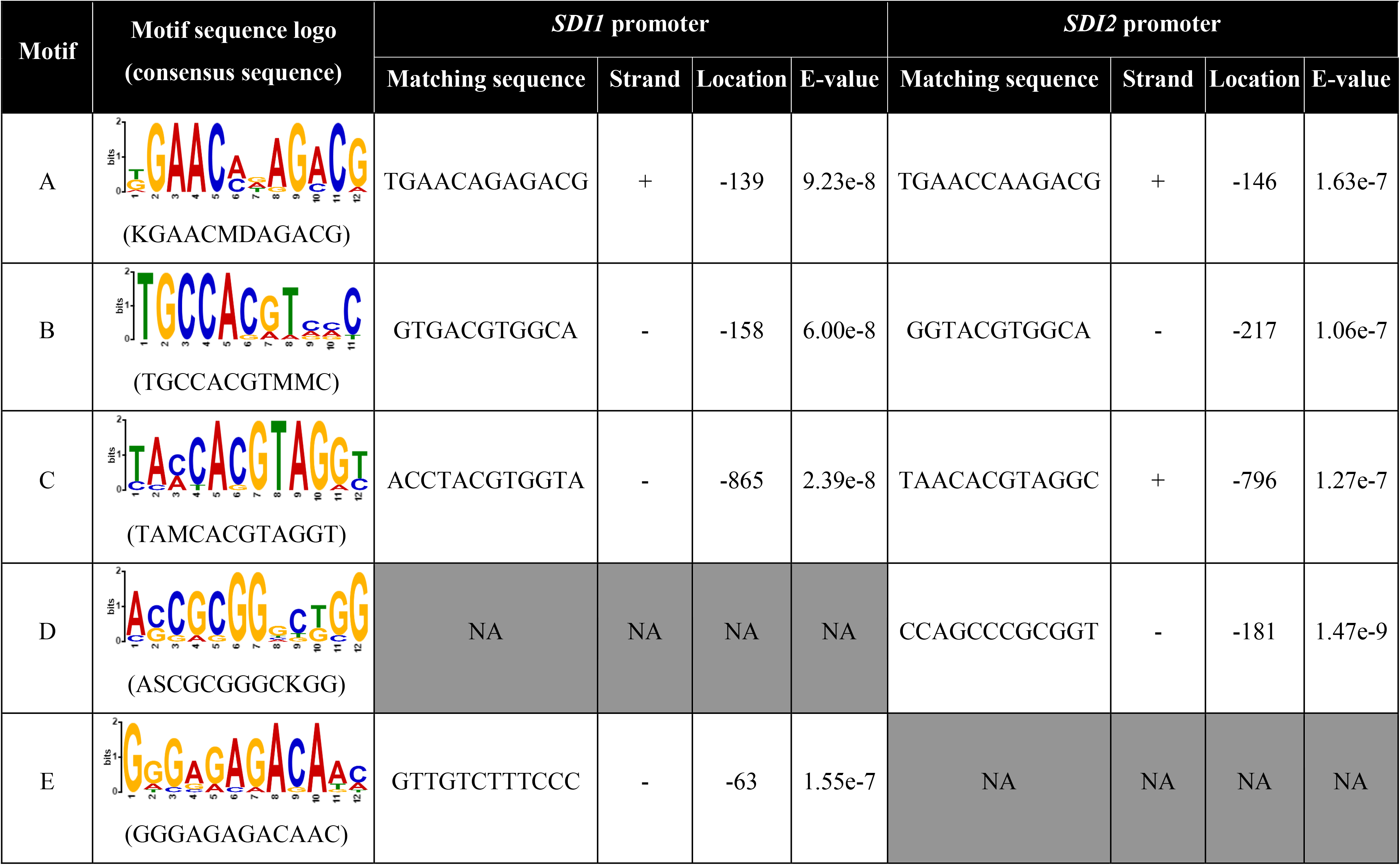
Sequence motifs predicted by MEME in *SDI1* and *SDI2* promoters. Five putative motifs identified by MEME-based comparative alignment of promoters of SDI-homolog genes in Brassicaceae are presented here. Ambiguous bases in each motif are represented with their IUPAC code: D stands for either G, A or T; K stands for either G or T; M stands for either A or C; and S stands for either C or G. The precise sequence in the *SDI1* and/or *SDI2* promoters matching the motif sequence is shown along with the match location, the match strand (sense strand (+); anti-sense (−)), and the sequence similarity (E-value) as determined by Blastp.

| Motif | Motif sequence logo<br>(consensus sequence) | SDI1 promoter |  |  |  | SDI2 promoter |  |  |  |
| --- | --- | --- | --- | --- | --- | --- | --- | --- | --- |
|  |  | Matching sequence | Strand | Location | E-value | Matching sequence | Strand | Location | E-value |
| A | (KGAACMDAGACG) | TGAACAGAGACG | + | -139 | 9.23e-8 | TGAACCAAGACG | + | -146 | 1.63e-7 |
| B | (TGCCACGTMMC) | GTGACGTGGCA | - | -158 | 6.00e-8 | GGTACGTGGCA | - | -217 | 1.06e-7 |
| C | (TAMCACGTAGGT) | ACCTACGTGGTA | - | -865 | 2.39e-8 | TAACACGTAGGC | + | -796 | 1.27e-7 |
| D | (ASCGCGGGCKGG) | NA | NA | NA | NA | CCAGCCCGCGGT | - | -181 | 1.47e-9 |
| E | (GGGAGAGACAAC) | GTTGTCTTTCCC | - | -63 | 1.55e-7 | NA | NA | NA | NA |

In order to identify potential similarities between the novel motifs identified with MEME and previously described *cis*-elements and their associated biological functions, we utilized the PLACE database. Notably, motif A contains the SURE *cis*-element (GAGAC) and shares high sequence similarity with the UPE box (AG[G/A]T[T/A]CATTGAA[T/C]CT [A/G]GAC[A/G]), both of which have been linked to transcriptional regulation of sulfur responsive genes (Maruyama-Nakashita et al., 2005; Wawrzyńska et al., 2010), as well as a putative SLIM1-binding motif ([T/A]G[T/A]A[T/C]C[T/A] [A/G]GAC[A/G]) sequence (O’Malley et al., 2016). Motif B contains a general bZIP TF family binding site (bZIP core *cis*-element, ACGT) (Jakoby et al., 2002), as well as sequence motifs that have been linked to response to light (GCCAC) (Hudson et al., 2003; Jiao et al., 2005), transcriptional regulation of ABI3-and ABA-responsive genes (ACGTG) (Nakashima et al., 2006), dehydration stress and dark-induced senescence (ACGT) (Simpson et al., 2003), and unfolded protein responses (CCACGTCA) (Martínez et al., 2003; Oh et al., 2003). Like motif B, motif C also contains the ACGTG sequence motif involved in transcriptional regulation of ABI3- and ABA-responsive genes (Nakashima et al., 2006). Motif D contains the calmodulin-binding/CGCG box (Yang et al., 2002), which is relevant to several biotic (Galon et al., 2008; Vadassery et al., 2012) and abiotic (Virdi et al., 2015) stress signaling cascades in plants. Motif E contains the auxin response factor (ARF) binding site (TGTCTC) (Goda et al., 2004) and the SURE *cis*-element (Maruyama-Nakashita et al., 2005).

As a final step of the *in silico* promoter analysis, FootprintDB was used to predict TFs that may bind to motifs A-E in the promoters of *AtSDI1* and *AtSDI2* (Table S3). Motif A was predicted to serve as a binding site for ANAC2 (ATAF1), ARR11, EIN3, GAL1, GAL3, GT-1, HAT3.1, NAC68, REF6 (EIN6), and SLIM1. TFs predicted to bind motif B include bZIP family TFs, namely bZIP3, bZIP16, bZIP42, bZIP44, bZIP53, GBF1 (bZIP41), GBF5 (bZIP2), and HYH (bZIP64). Motif C may serve as a binding site for ABI5 (bZIP39), ANAC55, HY5 (bZIP56), MYC2, PIF3, TOE1, and TOE2. TFs predicted to bind to motif D are in the AP2-EREBP TF family, such as ERF4, ERF6, ERF15, and ERF105. Motif E might serve as a binding site for ARF1, ANAC2, BPC1, BPC6, FRS9, WRKY23, and WRKY30. Notably, some TFs predicted by the FootprintDB to bind the novel motifs are associated with sulfur metabolism, namely ANAC2, ARF1, EIN3, HY5, SLIM1 (Watanabe and Hoefgen, 2019). Whereas the rest of the predicted TFs have not yet been reported to be associated with regulation of sulfur-related pathways and may be involved in *SDI* regulation in other conditions.

### Y1H screen to identify TFs able to bind the *SDI* promoters

We employed Y1H analysis to screen a library of 1395 Arabidopsis TFs (Castrillo et al., 2011) for binding to the promoters of *SDI1* and *SDI2*. Prior to screening, the autoactivation level was determined for the promoter sequences that would function as bait for TF binding. The full-length *SDI1* promoter (upstream region −1985 to 0 from ATG) was found to have some autoactivation activity, but it could be suppressed with 2 mM 3-aminotriazole (3-AT), whereas the autoactivation activity of the full-length *SDI2* promoter (upstream region −2085 to 0 from ATG) was still significant even with a high concentration (60 mM) of 3-AT (Figure S2). In order to achieve lower autoactivation activity, shorter fragments of the *SDI2* promoter were used as DNA-bait, and for comparability reasons also for *SDI1*.

The proximal (*prox*) promoter regions of *SDI1* and *SDI2* contain sulfur responsive *cis*-elements (SURE, UPE-like motif, and TEBS), bZIP binding sites (bZIP core and E box), LFY consensus and RAV1-A *cis*-elements, as well as novel putative motifs (A, B, D, and E) (Figure 1A and 1C). We therefore chose the *prox* promoter regions of both *SDI1* and *SDI2* as baits for Y1H screening. Farther upstream, the *SDI2* promoter is dense in TFBS, including MYC, E-box, bZIP core, and DPBF1&2 BS *cis*-elements (Figure 1A). Notably, most of the *cis*-elements we identified in the *SDI2* promoter (Table 1) are classified as binding sites for bZIP family TFs and are enriched in the *dist* promoter region of the SDI2 gene, but not in the *SDI1* promoter region at similar distance from the translation start site (Figure 1A).

For the Y1H screening, the *prox* region of the *SDI1* promoter was defined as the fragment from −263 to 0 from ATG (including the 60-bp 5’UTR), whereas the *SDI2 prox* promoter was defined as spanning from −307 to 0 from ATG (including the 82-bp 5’UTR). The *dist* region of the *SDI1* promoter was defined as being between −1656 and −1256 from ATG and the *SDI2 dist* promoter as spanning from −1682 to −1282 from ATG. These *prox* and *dist* promoter fragments of *SDI1* and *SDI2* were confirmed to have low autoactivation activity (Figure S2) and were used as baits in the Y1H analysis together with the full-length *SDI1* promoter to screen a library of 1395 Arabidopsis TFs for binding interactions (Castrillo et al., 2011) (Figure 2).

**Figure 2:**
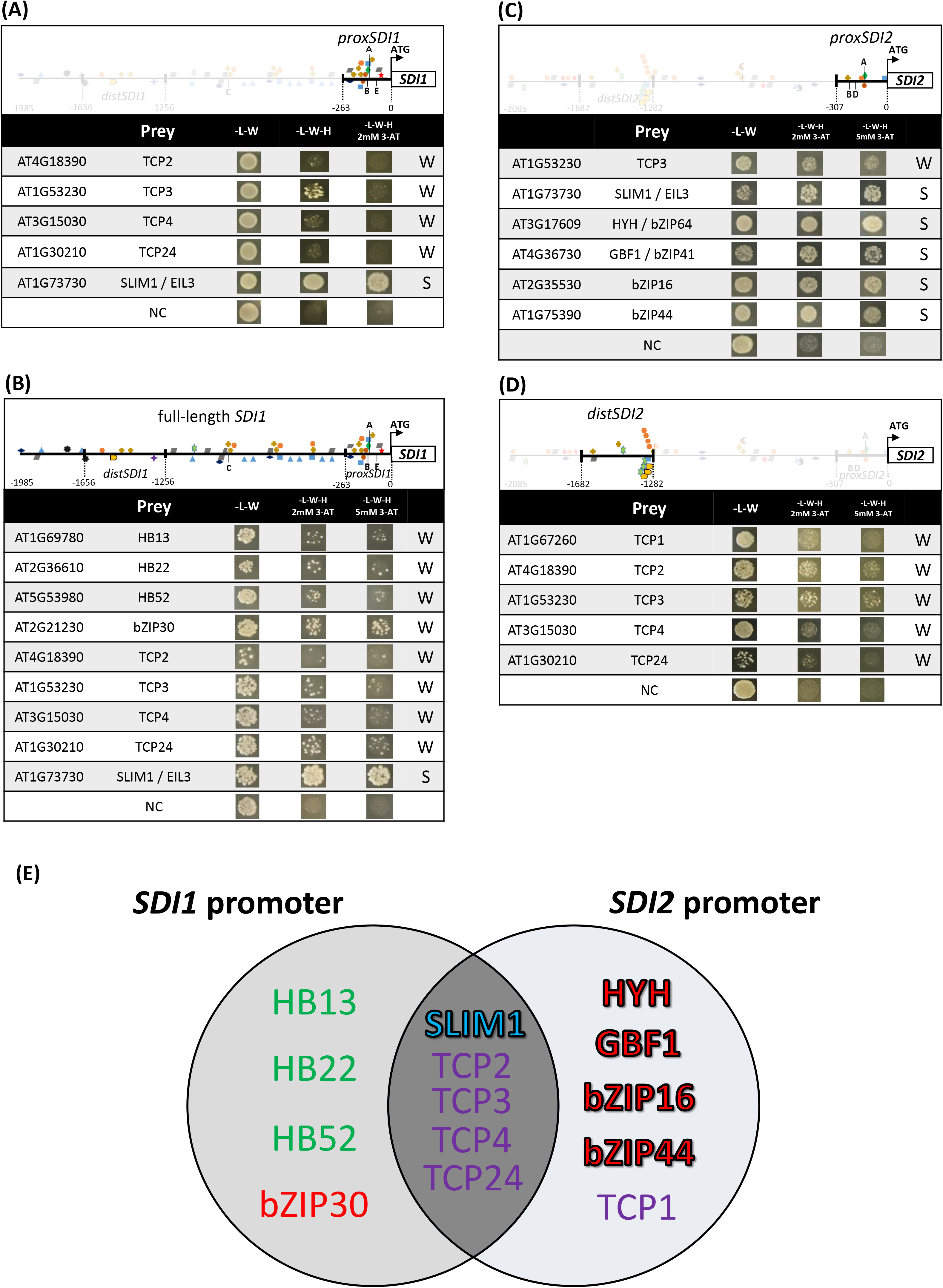
Y1H screening of Arabidopsis transcription factors for binding to *SDI1* and *SDI2* promoters. Selected regions of the *SDI1* (**A**, **B**) and *SDI2* (**C**, **D**) promoters were fused to the *HIS3* reporter gene and used as bait in Y1H screening of an Arabidopsis TF library. The cis-elements and putative motifs present in these promoter regions are represented with the same symbols and single letter codes as in Figure 1. Yeast were grown at 28°C for 4 days on media lacking leucine and tryptophan (-L-W) for the internal control and on media lacking leucine, tryptophan, and histidine (-L-W-H) including two different doses of 3-aminotriazole (3-AT) as a competitive inhibitor of the HIS3 enzyme to screen for binding. Negative control (NC) between bait and prey is also presented. The binding-strength between the TF and the bait region was classified as either strong (S) or weak (W) based on the extent of growth on –L-W-H media with 3-AT. Y1H screening results are summarized in a venn diagram (**E**). TFs are colored based on their TF family (blue for EIL, green for HB, red for bZIP, and purple for TCP). Strong TF-promoter Y1H interactions are indicated with bold font.

In total 14 TFs were identified in our Y1H analysis. Nine different TFs were found to bind the *SDI1* promoter bait sequences (Figure 2A, B). Five bound to the *SDI1 prox* promoter (Figure 2A), namely TCP2 (At4g18390), TCP3 (At1g53230), TCP4 (At3g15030), TCP24 (At1g30210), and SLIM1 (At1g73730). None bound the *dist* region of the *SDI1* promoter (data not shown). Nine TFs bound to the full-length *SDI1* promoter bait, including all five of the TFs that bound the *SDI1 prox* promoter fragment and four others, namely bZIP30 (At2g21230), HB13 (At1g69780), HB22 (At2g36610), and HB52 (At5g53980). Since these last four TFs were not found using the *prox* or *dist* promoter regions as bait, we infer that they bind to the promoter sequence between −1985 and −1656 and/or between −1256 and −263 from ATG. 10 TFs bound to *SDI2* promoter bait sequences: one to both the *dist* and *prox* regions, TCP3 (Figure 2C and 2D), five specifically to the *prox* region (Figure 2C), namely SLIM1, HYH (At3g17609), GBF1 (At4g3670), bZIP16 (At2g35530), and bZIP44 (At1g75390), and four specifically to the *dist* region (Figure 2D), namely TCP1 (At1g67260), TCP2, TCP4, and TCP24. By varying the concentration of 3-AT, we were able to tentatively classify the identified TFs as relatively strong binders (SLIM1, HYH, GBF1, bZIP16 and bZIP44) and weaker binders (bZIP30, HB13, HB22, HB52, TCP1, TCP2, TCP3, TCP4 and TCP24). Five TFs, namely SLIM1, TCP2, TCP3, TCP4, and TCP24, were found to interact with the promoters of both *SDI1* and *SDI2* (Figure 2E) with approximately equal binding strength.

### *In vitro* interaction between TFs and promoter fragments by EMSA

Since all TFs classified as strong binders by Y1H analysis were identified using the *prox SDI1* and *SDI2* promoters as bait (Figure 2A, C), we focused on these regions to validate interactions between the five strongly binding TFs and specific *cis*-elements. These *prox* promoter regions contain several known *cis*-elements, including bZIP core, SURE, TEBS, UPE-like motif, and E box, among others (Figure 3A). Short fragments from the *SDI1* and *SDI2 prox* promoter regions containing putative *cis*-elements for the strong binding TFs were chosen as probes for the EMSA analyses. Fluorescently labeled probes were used to detect interactions between a TF and the promoter fragment. Unlabeled competitor probes with and without mutated *cis*-element sequences were used to validate the specificity of the TF-DNA binding interaction.

**Figure 3:**
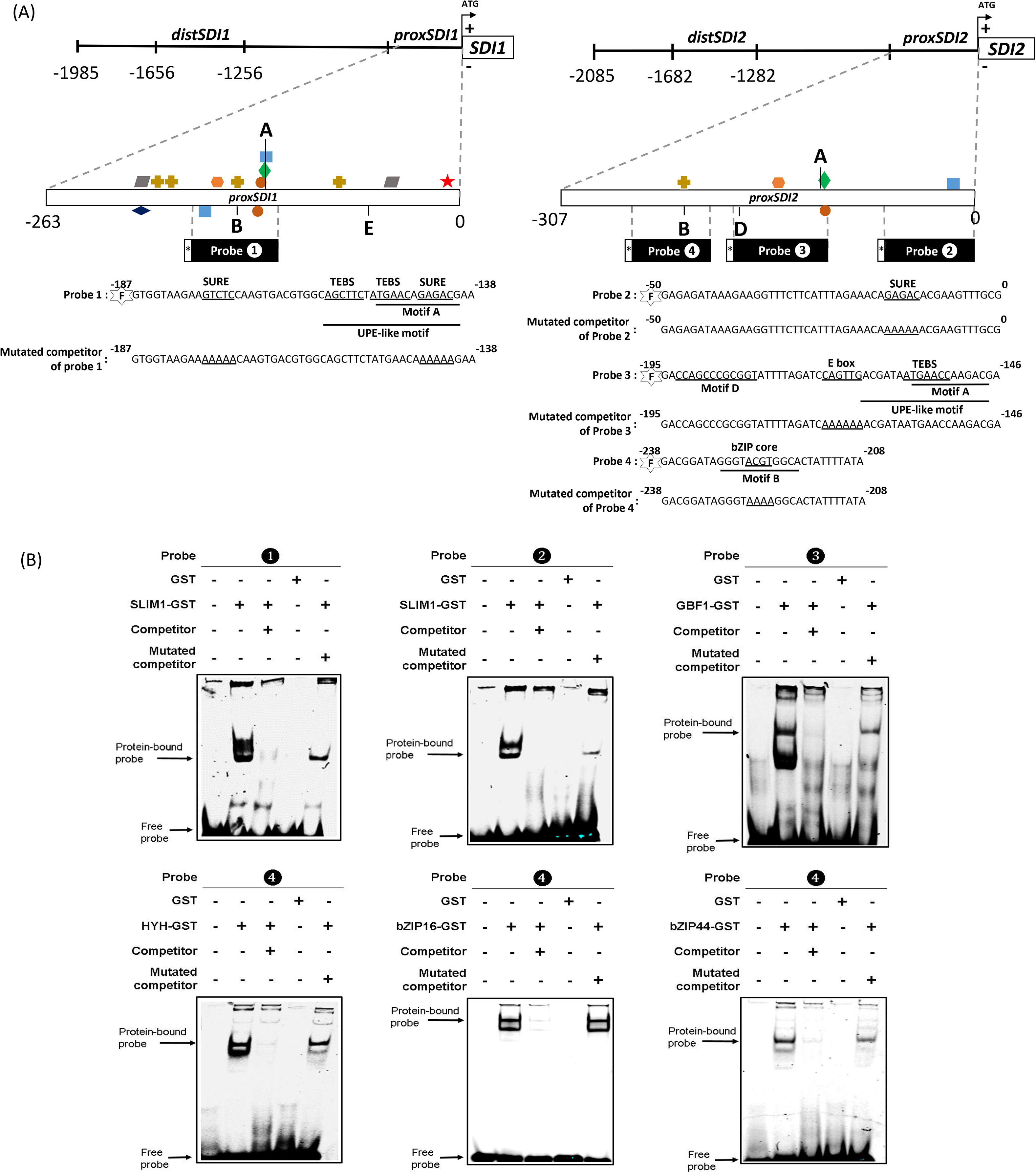
*In vitro* interactions between TFs and cis-elements in *SDI1* and *SDI2* proximal promoters. (**A**) Diagram of cis-elements and putative motifs in the proximal promoter regions of *SDI1* and *SDI2*. Symbols are the same as those used in Figure 1. The position and sequence of short probes from the *prox* regions of *SDI1* (probe 1) and *SDI2* promoters (probe 2, 3, and 4) are shown. Probes were fluorescently tagged with DY-682 (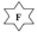) at their 5′ end. Competitor and mutated competitor probes were not fluorescently tagged. (**B**) Electrophoretic mobility shift assay results: SLIM1-binding activity to 50-bp DNA fragments containing the SURE elements from *proxSDI1* (probe 1) and *proxSDI2* (probe 2) promoters; GBF1-binding activity to a 50-bp DNA fragment containing the E box motif from the *proxSDI2* promoter (probe 3); Independent binding activities of HYH, bZIP16, and bZIP44 to a 30-bp DNA fragment containing the bZIP core motif from the *proxSDI2* promoter (probe 4). Shown are the 6% agarose gels used to analyze binding reactions of the indicated composition, where minus (−) indicates omission and plus (+) indicates addition.

SLIM1 was the only one of the five identified strong binders that bound to the *prox* promoter regions of both *SDI1* and *SDI2* in our Y1H screen (Figure 2A, C). SLIM1 is known to bind *in vivo* to TEBS *cis*-elements and UPE boxes found in the promoters of several -S inducible genes (Wawrzyńska et al., 2010). However, the promoters of some SLIM1-dependent sulfur deficiency-responsive genes and/or putative SLIM1-target genes contain neither a TEBS element, a UPE box, nor a UPE-like motif (Table S4), suggesting that SLIM1 might bind to other *cis*-elements as well. Interestingly, the promoters of most sulfur deficiency-responsive genes and putative SLIM1-targeted genes, including *SDI1* and *SDI2*, contain the SURE (GAGAC) sequence (Maruyama◻Nakashita et al., 2005; O’Malley et al., 2016). While most sulfur deficiency-responsive genes in roots are regulated by SLIM1 (Dietzen et al., 2020), SLIM1 has to the best of our knowledge not yet been shown to bind to the SURE element. The *prox* promoter of *SDI1* contains two SURE elements: one near the UPE-like motif and the other within it. In contrast, the UPE-like motif and the single SURE element in the *prox* promoter of *SDI2* are more distant from each other (Figure 3A). In order to assess whether the SURE *cis*-element is involved in SLIM1 binding to the *prox* promoter regions of *SDI1* and *SDI2*, we performed EMSA (Figure 3B, probe 1 and 2). We found that the SLIM1 protein bound to both probe 1 and 2. The intensity of the shifted band was very strongly decreased when unlabeled competitor probe was included in the binding reaction. In contrast, the shifted band was clearly visible when the SURE *cis*-element sequence(s) in the unlabeled competitor probes were modified to AAAAA. This indicates that SLIM1 can bind to the SURE element(s) of *SDI1* and *SDI2* promoters.

The other four strongly binding TFs, GBF1, HYH, bZIP16, and bZIP44, are members of the bZIP TF family and were identified by Y1H with the *prox* promoter of *SDI2* as bait. GBF1 has previously been reported to bind to the E box *cis*-element (CANNTG) (Maurya et al., 2015), which is present in the *prox* region of the *SDI2* promoter. Therefore, we performed EMSA with GBF1 and a *SDI2 prox* promoter probe containing the E box element (Figure 3B, probe 3). GBF1 was shown to bind to probe 3 and inclusion of an unlabeled probe 3 in the binding reaction resulted in strongly decreased intensity of the shifted band. Inclusion of an unlabeled competitor probe with the E box mutated to AAAAAA moderately reduced the intensity of the shifted band. We conclude that GBF1 can bind directly to the *SDI2* promoter via interaction with the E box *cis*-element in the *prox* region. Our *in silico* FootprintDB analysis (Table S3) predicted that the other three strongly binding TFs, namely bZIP16, bZIP44, and HYH (bZIP64), can bind to motif B, which contains a bZIP core sequence (ACGT). We tested by EMSA whether bZIP16, bZIP44, and HYH can bind directly to a *SDI2 prox* promoter probe containing the bZIP core element within motif B (Figure 3B, probe 4). We observed binding of these three bZIP TFs to probe 4 and that the presence of unlabeled competitor probe strongly reduced the intensity of the shifted band. In addition, inclusion of an unlabeled competitor with a mutated bZIP core did not significantly decrease the intensity of the shifted band. Thus, the single bZIP core element in the *SDI2 prox* promoter is likely to be a binding site for bZIP16, bZIP44, and HYH. It remains to be investigated how these three TFs may compete and/or cooperate for binding to this *cis*-element *in planta*.

### Transcriptional response of candidate regulators of *SDIs* under conditions which alter *SDI* expression

Since expression of *SDI1* and *SDI2* are known to respond strongly to S-deficiency, we tested whether expression of the putative TF-regulators of *SDI* from the Y1H screen are also sensitive to sulfur status. Arabidopsis seedlings were grown in shaking cultures under full nutrient (FN) conditions for eight days and then shifted to sulfate free media (-S) or to fresh FN media. Four days after the shift, the sulfate level in -S seedlings was approximately 25% of the level in FN seedlings (Table S5). As expected, *SDI1* and *SDI2* expression was highly induced by withdrawal of sulfate from the media, consistent with several published microarray experiments from the GEO and ArrayExpress databases (Figure 4; Table S5). However, neither in our shaking culture experiments nor in the publicly available microarray studies did we find clear indication that transcript levels of the 14 TF candidates are responsive to sulfate deficiency.

**Figure 4:**
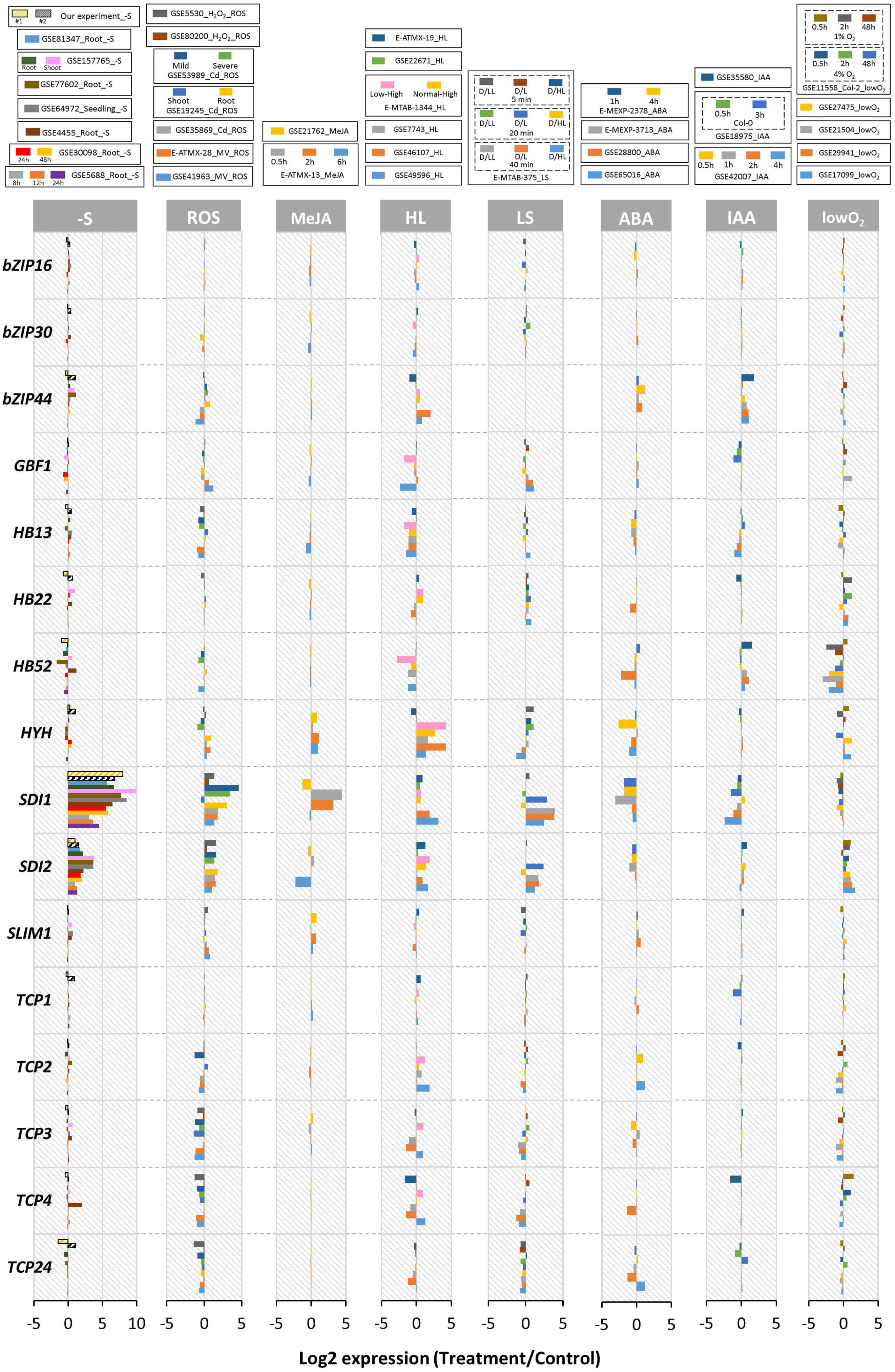
Transcriptional response of candidate regulators of *SDI*s under conditions which alter *SDI* expression. Meta-analysis of log2 fold change (treated versus control) of 14 TF candidates under 8 different conditions that alter *SDI* expression: sulfur starvation (-S) including our experiments in seedlings (#1 and #2), oxidative stress (ROS), exogenous application of methyl jasmonic acid (MeJA), high light (HL), light-shift (LS), exogenous application of abscisic acid (ABA) or auxin/indole-3-acetic acid (IAA), and hypoxia (lowO_2_), based on published transcript profiling data from the GEO and ArrayExpress databases.

We then asked whether the expression levels of these 14 TFs are regulated by other conditions that alter expression of *SDI1* and/or *SDI2* (Figure 4, Table S5). A fairly consistent expression change across datasets was evident for several of the candidate TFs: high light stress tends to induce *HYH* and repress *HB13* expression, while hypoxia tends to repress *HB52*. Notably, four of the five TFs that were found by Y1H to strongly bind the promoters of *SDI*s, do not show clear expression changes in conditions that alter *SDI* expression. This includes *SLIM1*, despite it being the only validated TF regulator of *SDI1* and *SDI2* induction under sulfur deficiency (Maruyama-Nakashita et al., 2006; Dietzen et al., 2020). Clearly, the lack of transcriptional response of a TF to a particular condition does not rule out an important role for that TF in regulation of *SDI1* and/or *SDI2* in that condition.

## DISCUSSION

The primary goal of this study was to identify TFs and promoter sequence elements that may participate in transcriptional regulation of *SDI1* and *SDI2*. To this end we screened a large library of TFs by Y1H for binding to *SDI* promoters. The Y1H screen identified 14 TFs belonging to four TF classes based on PlnTFDB 3.0: bZIP, EIL, HB, and TCP TF families. Five TFs were classified by the Y1H screen as strong binders: the EIL family member SLIM1/EIL3, which bound to both *SDI1* and *SDI2* promoters, and four bZIP family TFs, namely bZIP16, bZIP44, GBF1, and HYH, all of which bound to *SDI2* promoter fragments but not detectably to the promoter of *SDI1*. The three HB family and five TCP family TFs appeared to bind only weakly to the *SDI* promoter regions tested by Y1H. The number of TFs found by Y1H screening to interact with the *SDI* promoters is less than might be predicted from our *in silico* analysis. There are many potential sources of false-negatives in Y1H screens, including toxicity of the strongly expressed Arabidopsis TF-GAL4 activation domain fusion protein, or conversely, poor nuclear-targeting or protein stability of the prey protein (Liu et al., 1993). TF-DNA binding can also be strongly affected by growth condition-dependent post-translational modifications that are not present in the *Saccharomyces cerevisiae* cultures used for Y1H (Walhout, 2011). And to facilitate binding to DNA certain TFs depend on physical association with other protein factors, which may not be present in yeast cells (Hickman et al., 2013). Further, transcriptional activation in *Saccharomyces cerevisiae* tends to decline as the distance between the binding element and the transcriptional start site increases (Dobi et al., 2007). SDI1 and SDI2 were included in the Y1H prey library because *SDI1* and *SDI2* had previously been shown to negatively regulate the expression of each other (Aarabi et al., 2016). Neither SDI1 nor SDI2 were among the 14 TFs identified in our Y1H screening (data not shown). SDI proteins bind through their TPR domain to other proteins, as we have previously shown for MYB28 (Aarabi et al., 2016). Therefore, we conclude that the mutual negative regulation of *SDI1* and *SDI2* might be exerted through interaction with other TFs.

### SLIM1 binds to SURE elements in *SDI* promoters

Among the TFs identified by Y1H, only SLIM1 bound strongly to both the *SDI1* and *SDI2* promoters. SLIM1 has previously been implicated in transcriptional responses to sulfur deprivation (Maruyama-Nakashita et al., 2006; Dietzen et al., 2020), a condition that strongly induces expression of *SDI*s (Howarth et al., 2009; Aarabi et al., 2016) (Figure 4). The *SDI1* and *SDI2* proximal promoter regions to which SLIM1 strongly bound in our Y1H screen contain several sequence elements that we suspected may be important for binding. SLIM1 has previously been shown to bind the UPE box and TEBS *cis*-element *in vivo* (Lewandowska et al., 2010; Wawrzyńska et al., 2010). The sulfate response element SURE is also present in the proximal promoter regions of *SDI*s. Further, by comparing *SDI* promoters of various *Brassicaceae* species we identified a conserved motif, termed motif A ([T/G]GAAC[A/C][A/G/T]AGACG), which contains the SURE element (GAGAC) (Figure 1B, Table 2). Notably, motif A is highly similar to a motif enriched in SLIM1 binding sites identified by DAP-seq ([T/A]G[T/A]A[T/C]C[T/A][A/G]GAC[A/G]) (O’Malley et al., 2016). 50-bp regions of the proximal promoters containing motif A were used as probes in EMSA analyses, and we demonstrated that SLIM1 can bind to these promoter fragments *in vitro*. Unlabeled competitor probes in which the SURE elements were mutated were weaker competitors of SLIM1 binding than unmutated, unlabeled probes. This indicates that the SURE element(s) in the proximal promoters of *SDI1* and *SDI2* likely contribute substantively to binding of SLIM1 to these promoters (Figure 3). To the best of our knowledge this is the first time SLIM1 binding to the SURE *cis*-element has been shown, and this finding is of particular importance because of the major role SLIM1 has in regulating sulfate deprivation responses (Maruyama-Nakashita et al., 2006).

Interestingly, the SURE elements in the *proxSDI1* and *proxSDI2* promoter regions are either adjacent to or embedded in palindromic sequences (Figure S3). Palindromic sequences in double stranded DNA can form stable cruciform structures *in vivo* (Brázda et al., 2011; Gentry and Hennig, 2016), and there is some evidence that they may influence the specificity and/or affinity of certain TFs, including ethylene-insensitive3 (EIN3) and EIN3-like (EIL) TFs (Yamasaki et al., 2005). The *distSDI2* promoter region also contains a SURE element, but it is not located near a palindromic sequence, and when this region was used as bait in our Y1H, it did not result in SLIM1 binding. This is consistent with Y1H screening by Wawrzyńska et al. (2010) in which a bait sequence containing the SURE element without any adjacent palindrome was incapable of binding to NtEIL2 (a functional orthologue of AtSLIM1).

To investigate the prevalence of potential SLIM1 binding sites (UPE box, UPE-like motif, TEBS, SURE, and motif A) we performed *in silico* analysis on promoters of 84 –S responsive SLIM1-dependent genes (Maruyama-Nakashita et al., 2006) and 184 promoters of genes bound by SLIM1 in DAP-seq analysis (O’Malley et al., 2016) (Table S4). Interestingly, approximately one third of the promoters in each gene list contain SURE but not one of the SLIM1 binding sites identified previously in literature, namely UPE-box, UPE-like box, and TEBS. For two of these genes, *BGLU28* (*At2g44460*) and *HAL2-LIKE* (*At5g54390*), we searched for palindromic sequences located near the SURE sites in their promoters. These genes are of particular interest because in addition to lacking a UPE-box, UPE-like, and TEBS, they were identified as putative SLIM1-target genes by both DAP-seq (O’Malley et al., 2016) and transcriptome analysis of *slim1* mutants under sulfur starvation (Maruyama-Nakashita et al., 2006). In both promoters, the SURE element most proximal to the translation start site is adjacent to a palindromic sequence (Table S6). Taken together, these analyses suggest that the proximity of palindromic sequences to a SURE element might help explain SLIM1-dependent transcriptional responses of genes lacking either a UPE box or UPE-like motif and/or TEBS in their promoter. Bioinformatic and *in vitro* binding studies will be necessary to validate whether, and if so how, cruciform structures and the precise location of the palindromic sequences relative to SURE affect SLIM1 binding and thus contribute to the complex regulation of the SDI genes.

### *SDI1* and *SDI2* responses in conditions other than S deficiency

Although expression of both *SDI1* and *SDI2* respond to sulfur deficiency, induction of *SDI1* in these conditions is far stronger than induction of *SDI2* (Figure 4) (Maruyama-Nakashita et al., 2006; Aarabi et al, 2016; Dietzen et al., 2020). Similar differential induction strength is observed with selenate treatment, which is a mimic of sulfate starvation (Van Hoewyk et al., 2008 GEO: GSE9311), and with application of cadmium and menadione, which generate oxidative stress (López-Martín et al., 2008 ArrayExpress: E-GEOD-19245; Jobe et al., 2012 ArrayExpress: E-GEOD-35869; Lehmann et al., 2009) (Figure 4). In contrast, hypoxic conditions have a stronger effect on *SDI2* expression than on *SDI1* (Christianson et al., 2010 GEO: GSE21504; Licausi et al., 2010 GEO: GSE17099; Gibbs et al., 2011 GEO: GSE29941) (Figure 4). Further, exogenous application of phytohormones such as abscisic acid and auxin results in downregulation of *SDI1* but little change in *SDI2* expression (Mizoguchi et al., 2010 ArrayExpress: E-MEXP-2378; Kim et al., 2011 GEO: GSE28800; Bargmann et al., 2013 GEO: GSE35580; Lewis et al., 2013 GEO: GSE42007; Kim et al., 2019 GEO: GSE65016) (Figure 4). Taken together these results point to differential molecular regulation of *SDI1* and *SDI2* expression under some conditions.

12 of the 13 TFBS that we localized to the *SDI* promoters with database and literature searches are found in potential regulatory regions of both *SDI1* and *SDI2* (Table 1). However, several binding sites are found predominantly in the promoter of one homolog. There are nine W-boxes, a WRKY TF family binding site, in the region 2000-bp upstream of the translation start site of *SDI1*, but only one in the same promoter region of *SDI2*. Similarly, the bZIP core binding site is present in 11 and 6 copies in the promoters of *SDI1* and *SDI2*, respectively. In contrast, two other binding sites for bZIP family TFs, the DPBF1&2 and E box, are significantly more prevalent in the promoter of *SDI2* than *SDI1*.

In addition to the unequal prevalence of particular *cis*-elements in the two promoters, our Y1H results suggest that various TFs may have higher affinity for the promoter of one *SDI* homolog than the other (Figure 2). For example, four bZIP family TFs, bZIP16, bZIP44, GBF1, and HYH, were classified as strong binders to the *SDI2* promoter, but binding to the *SDI1* promoter was not detectable by Y1H. bZIP16, GBF1, and HYH are involved in regulation of light-mediated processes in plants (Hsieh et al., 2012; Mallappa et al., 2006; Holm et al., 2002). These TFs may be involved in *SDI2* induction by high light stress as well as by shifting plants from light to dark (Figure 4). bZIP44 expression is important during early steps in germination (Iglesias◻Fernández et al., 2013), and notably, SDI2 induction is strongly induced at a similar stage (Narsai et al., 2011 GEO: GSE30223). Given our Y1H and EMSA results, it is plausible that bZIP44 may positively regulate *SDI2* expression during seed germination. However, given that multiple bZIP TF binding sites are found in the promoters of both *SDI* homologs, it is not straightforward to explain the preference of bZIP16, bZIP44, GBF1, and HYH for *SDI2* in our Y1H. Although the proximal region of both *SDI1* and *SDI2* promoters contain an E box, a palindromic sequence is only observed near the E box of *proxSDI1*. The precise sequence context of the bZIP binding site (Williams et al., 1992; Fujii et al., 2000) and perhaps genomic DNA secondary structures (Fujii et al., 2000) may be important for differential binding of these TFs to the promoters of *SDI1* and *SDI2*. If differential association also occurs *in vivo*, it could contribute to a molecular basis for differential expression of *SDI1* and *SDI2*.

In conclusion, previously published work has shown that *SDI1* and *SDI2* are important for S-metabolism by reducing GSL biosynthesis when sulfate availability is low, and in this context at least, the homologs are able to complement each other’s function (Aarabi et al., 2016). However, our meta-analysis clearly suggests that *SDI1* and *SDI2* have functions in contexts beyond S-deficiency and that they are likely to have some non-redundant functions. What these functions may be and the nature of the metabolic and signaling pathways involved remain open questions. Here we have shown SLIM1, GBF1, HYH, bZIP16, and bZIP44 binding to *SDI* promoter sequences by Y1H and EMSA. And with the exception of SLIM1, these TFs have not previously been associated with S-deficiency responses generally or *SDI* regulation more specifically. While we cannot yet conclude that these TFs are important for regulation of *SDI1* and/or *SDI2 in vivo*, their identification as candidate regulators is a clear step forward and opens the door to more targeted studies to address the molecular details of transcriptional regulation and biological function of SDI1 and SDI2.

## EXPERIMENTAL PROCEDURES

### *In silico* analysis

The AtcisDB (https://agris-knowledgebase.org/AtcisDB; Davuluri et al., 2003; Yilmaz et al., 2011), PLACE database (https://www.dna.affrc.go.jp/PLACE version 30.0, 2007 update; Higo et al., 1999), and literature reviews were used to identify putative *cis*-regulatory elements in the promoters of *AtSDI1* (*At5g48850*) and *AtSDI2* (*At1g04770*). Full-length promoter sequences (approximately 2000-bp upstream from ATG) of SDI homologs in the Brassicaceae family identified by PLAZA version 4.0 (http://bioinformatics.psb.ugent.be/plaza; Van Bel et al., 2018) were used as input for *in silico* detection of novel putative motifs. Multiple EM for Motif Elicitation (MEME) suite version 5.1.1 (http://meme-suite.org/tools/meme; Bailey et al., 2009) was used to identify conserved DNA sequence motifs and to visualize their localization in the promoters. Parameters for the motif search were the following: maximum of 20 unique motifs per sequence, motif length 10 – 12 nucleotides, and motif E-value ≤ 0.05, which is depended on its log likelihood ratio, motif length, frequencies of a motif appearance in the set construction. In order to predict TFs that may bind to the novel putative motifs identified by MEME, we used the FootprintDB (http://floresta.eead.csic.es/footprintdb; Sebastian et al., 2014; Contreras-Moreira et al., 2016) with an E-value cut off of 10^−3^, which is calculated by Blastp alignments against the 3D-footprint library. TFs were assigned to families according to the PlnTFDB 3.0 (http://plntfdb.bio.uni-potsdam.de/v3.0; Pérez-Rodríguez et al., 2010). Additionally, published transcript profiling datasets deposited in Gene Expression Omnibus (GEO) (http://www.ncbi.nlm.nih.gov/geo; Barrett et al., 2007) and ArrayExpress (https://www.ebi.ac.uk/arrayexpress; Athar et al., 2019) were used to assess transcript levels of TF candidates in response to various conditions. The palindromic sequences within the promoters of SDI genes were detected by DNA analyser (http://bioinformatics.ibp.cz; Brázda et al., 2016). Patmatch (https://www.arabidopsis.org/cgi-bin/patmatch/nph-patmatch.pl; Yan et al., 2005) was used to determine the presence of *cis*-elements and motifs of interest in the promoters of putative SLIM1-target genes (Table S4).

### Construct cloning

Bait constructs for Y1H analysis were produced by inserting promoter regions of SDI genes into the multiple cloning site of pTUY1H using In-Fusion^®^ HD cloning kit (Takara Bio Inc.) and then used to transform yeast strain Y187, mating type alpha. To produce proteins for EMSA, coding sequences without stop codons of candidate TFs were amplified via PCR from *A. thaliana* (Col-0) and then cloned into pENTR™/D-TOPO™ (Invitrogen) vector. Gateway recombinant cloning was performed to subclone the aforementioned constructs into pDEST™24 (Invitrogen) using a LR cloning reaction (Invitrogen). Destination constructs coded for C-terminal GST-tagged TF proteins for expression in *E. coli* rosetta (DE3).

### Yeast One-Hybrid (Y1H) Screening

A detailed description of the Y1H screening procedure can be found in Castrillo et al. (2011). Briefly, a library of 1395 *A. thaliana* TF fused to Gal4 activation domain within pDEST™22 (Invitrogen) constructs in *S. cerevisiae* strain YM4271, mating type a, served as prey for screening with the bait constructs described above (see construct cloning). DNA bait-TF prey interactions were identified on synthetic defined medium lacking Leu, Trp, and His. Two different concentrations of 3-amino-1,2,4-triazol (3-AT), 2 mM and 5 mM, were applied on the medium in order to roughly assess the binding-strength between candidate TFs and *SDI* promoter regions.

### TF protein expression and Western blot analysis

*E. coli* rosetta (DE3) cells were transformed with C-terminal GST tagged Arabidopsis TF expression constructs or GST (see above, construct cloning) and shaken (200 rpm) overnight at 37°C in 3 mL Luria-Bertani (LB) media with ampicillin (100 μg/ml) and chloramphenicol (50 μg/ml) in order to select a positive clone which contains pDEST™24 (Amp^R^) and pRARE (Cam^R^). In the morning, 3 mL fresh LB media with ampicillin and chloramphenicol was inoculated with 150 μL of the overnight pre-culture and shaken for 1.5 h. TF expression induction was initiated with 1mM isopropyl β-D-1-thiogalactopyranoside (IPTG) at 30 °C and allowed to proceed for 6h. 2 mL of the induced culture was collected, the cells pelleted by tabletop microcentrifugation, and the cell pellets resuspended in 150 μL extraction buffer [20 mM sodium-phosphate buffer (pH 7.4), 0.5 M NaCl, 1 mM phenylmethylsulfonyl fluoride (PMSF), 1mM ethylenediaminetetraacetic acid (EDTA), 5 μL protease inhibitor cocktail solution for use with bacterial cell (P8465, Sigma)]. Cell disruption was achieved by enzymatic and mechanical approaches, lysozyme and ultrasonication, respectively. The extracts (crude, supernatant, and pellet) for each target protein were collected independently. The presence of full-length TF proteins was confirmed by a western blot signal, via the 800 nm channel of the Odyssey^®^ 9120 (LI-COR), at the predicted molecular weight with a fluorochrome-conjugated secondary antibody (Anti-DCX antibody produced in goat, Sigma) against a primary antibody (GST-tag Monoclonal antibody, Novagen), which is attached to GST-tagged TF proteins (Figure S4).

### Electrophoretic Mobility Shift Assay (EMSA)

To create double-stranded probes, 10 μM each of complementary oligonucleotides (Table S1) from Eurofins Genomics (https://www.eurofinsgenomics.eu/) were mixed and melted at 95◻°C for 5◻min in TE buffer in a T100™ thermocycler (Bio-rad). Then the denatured oligonucleotides were allowed to anneal as the temperature was reduced by 1 °C every 20 sec, until reaching 4° C, to create double-stranded probes. Fluorescently labeled (5′ DY-682) dimerized probes were then diluted 1:300 in TE buffer. Unlabeled dimerized probes were not diluted. The binding reactions between the supernatant of candidate TFs extracted from transformed DE3 cells (see above, TF protein extraction and Western blot analysis) and dimerized probes were performed with the Odyssey^®^ Infrared EMSA kit (LI-COR) at room temperature in the dark followed by electrophoresis through a 6% DNA retardation gel (Invitrogen) run in TBE buffer at 4°C in the dark. In addition, the supernatant from a crude extract of *E. coli* transformed with the cloning entry vector was used as a control for GST binding to each DNA probe. Finally, the mobility of the labeled DNA probe was visualized via the Odyssey^®^ 9120 (LI-COR) in the 700 nm channel.

### Plant material and shaking culture conditions

*Arabidopsis thaliana* ecotype Col-0 seeds were sterilized with chloride gas for 3h and were put into 100 mL Erlenmeyer flasks containing 30 mL FN (0.75 mM MgSO_4_, 1.5 mM Ca(NO_3_)_2_, 0.1 mM FeEDTA, 0.75 mM KH_2_PO_4_, 1 mM KNO_3_, 0.1 μ CoCl_2_, 0.5 μM CuCl_2_, 50 μ H_3_BO_3_, 1 μ KI, 10 μ MnCl_2_, 0.8 μM Na_2_MoO_4_, and 1.75 μM ZnCl_2_) liquid medium and stratified for 2 days, at 4°C in the dark. Sterile, stratified seeds germinated and were grown in shaking culture (85 rpm) with 50 μE m^−2^ s^−1^ light intensity at 22°C for 8 days, then were transferred into flasks with fresh FN media or –S media (0 mM MgSO_4_) and allowed to grow in shaking culture for an additional 4 days. Seedlings from independent flasks were harvest and washed in deionized water before freezing in liquid nitrogen.

### Sulfate content analyzed by ion chromatography

50 mg ground samples of Arabidopsis seedlings were dissolved in the extract mixture (MTBE:MeOH, 3:1, vol/vol), and then vortexed and sonicated for 15 min. A fractionation by phase separation was performed via inclusion of H_2_O:MeOH (3:1, vol/vol) as well as centrifugation (14,000 rpm) for 5 min at 4°C. 100 μL polar phase supernatant was diluted into 550 μL UPLC-grade water. After vortexing, the sample was centrifuged at 14000 rpm at 4°C. Sulfate content in the supernatant was determined by high performance anion exchange chromatography with conductive detector using Dionex IC3000 (Dionex).

### qRT-PCR based analysis of candidate TF expression

Total RNA was extracted with TRIzol™ reagent (Invitrogen) from 100 mg ground samples of Arabidopsis seedlings. 2 units TURBO™ DNase (Invitrogen) was added to mitigate DNA contamination prior to first-strand cDNA synthesis with PrimeScript™ RT reagent kit (Takara Bio) from approximately 1 μg total RNA. Quantitative RT-PCR was performed with 7900 HT fast realtime PCR system (Applied Biosystems) using gene-specific primers of candidate TFs (Table S2) and fluorescent dye SYBR™ Green (Applied Biosystems). Expression values relative to *ACT2* were calculated with the comparative ΔΔCt method (Schmittgen and Livak, 2008).

## DATA AVAILABILITY STATEMENT

Any related details can be found in the manuscript and its supporting materials.

## ACKNOWLEDGMENTS

We would like to especially thank Dr. Salma Balazadeh and Dr. Arun Sampathkumar, Max-Planck Institute of Molecular Plant Physiology, for helpful discussions, as well as Karina Schulz (member of Balazadeh lab) for preparing the arrayed Arabidopsis TF library. We thank Dr. Dirk Walther, Max-Planck Institute of Molecular Plant Physiology, for suggesting bioinformatics tools, as well as Franziska Brückner (member of Hoefgen lab) for measuring sulfate content. AA has been funded by the Deutsche Forschungsgemeinschaft (DFG), HO1916/13-1. AR, SJW, and RH received institutional funding by the Max Planck Society.

## AUTHOR CONTRIBUTIONS

AR, SJW, and RH conceptualized and designed the research study. AR executed the bioinformatic analyses of promoter sequences as well as *in silico* meta-analysis of publicly available transcript profiling experiments. AR conducted the Y1H and EMSA experiments. AR and AA performed the qPCR analysis of 14 TFs plus SDI1 and SDI2 genes under normal and deficient sulfur supply. AR wrote the manuscript, assisted by all the authors, with particular support from SJW and RH.

## SUPPORTING MATERIALS

Figure S1: Moderate expression of *SDI1* and *SDI2* under various conditions in many developmental stages of Arabidopsis, based on the Genevestigator database

Figure S2: Examination of autoactivation of proximal-, distal-, and full-length regions of *SDI1* and *SDI2* promoters

Figure S3: Palindromic sequences within proximal regions of SDI1 and SDI2 promoters

Figure S4: Western blotting of the candidate TFs fused to the GST protein

Table S1: Probe sequences for EMSA

Table S2: List of qRT-PCR primers for validating transcript levels of TF candidates bound to *SDI1* and *SDI2* promoters

Table S3: List of TFs predicted by FootprintDB to bind to putative motifs A through E

Table S4: Motif and *cis*-element search results in promoters of putative SLIM1-target genes

Table S5: Transcript response data for Figure 4

Table S6: Sequence context of the most proximal SURE element in the promoters of two presumed SLIM1-target genes lacking a UPE box, UPE-like motif or TEBS in their promoters

