## Supplementary material for "*In silico* analysis of *cis*-elements and identification of transcription factors putatively involved in the regulation of the OAS cluster genes *SDI1* and *SDI2*": Figure S1

**Figure S1: Moderate expression of *SDI1* and *SDI2* under various conditions in many developmental stages of Arabidopsis, based on the Genevestigator database**

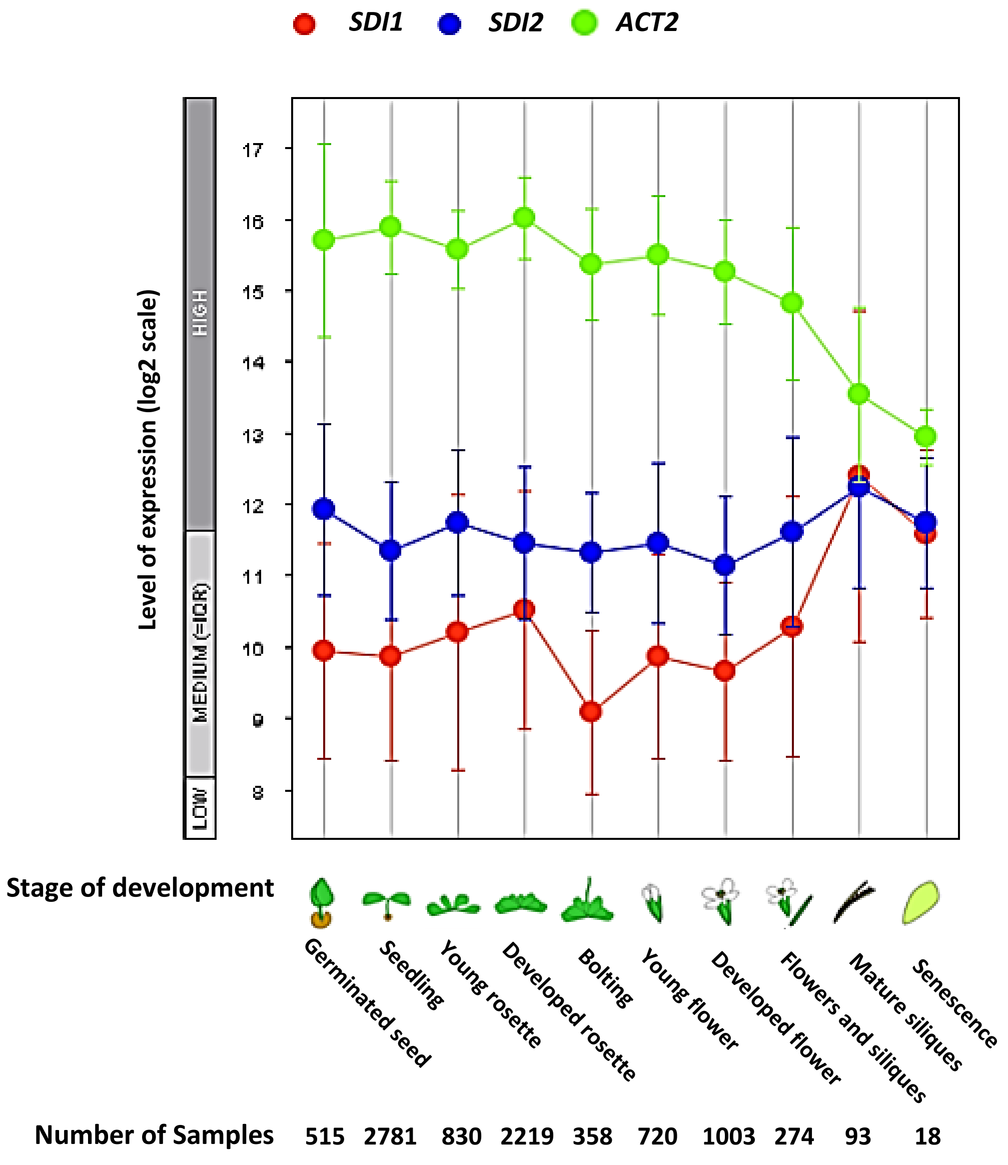
