## Supplementary material for "*In silico* analysis of *cis*-elements and identification of transcription factors putatively involved in the regulation of the OAS cluster genes *SDI1* and *SDI2*": Figure S2

**Figure S2:** Examination of autoactivation of proximal-, distal-, and full-length regions of *SDI1* and *SDI2* promoters

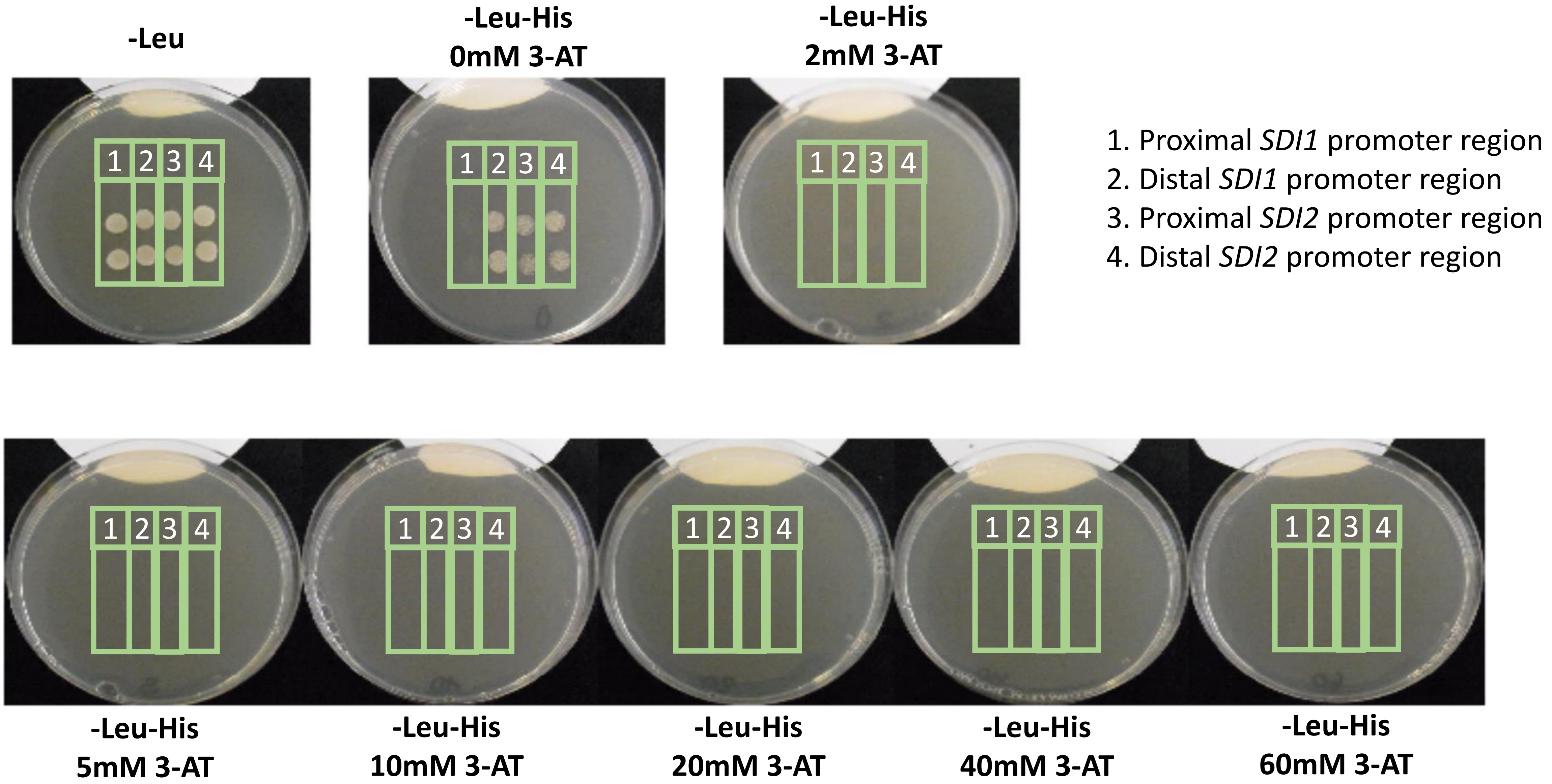

**Figure S2:** Examination of autoactivation of proximal-, distal-, and full-length regions of *SDI1* and *SDI2* promoters

5. Full-length *SDI1* promoter region
6. Full-length *SDI2* promoter region

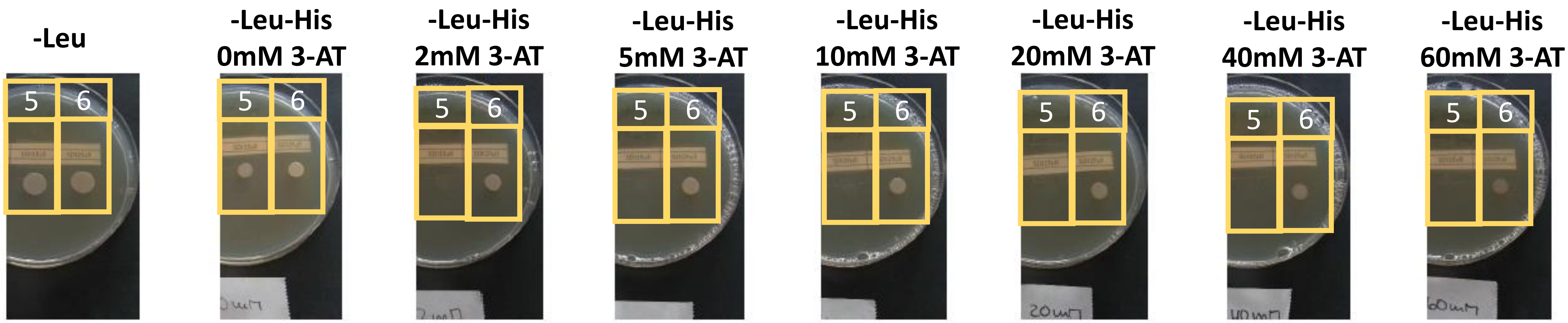
