## Supplementary material for "*In silico* analysis of *cis*-elements and identification of transcription factors putatively involved in the regulation of the OAS cluster genes *SDI1* and *SDI2*": Figure S3

**Figure S3:** Palindromic sequences within proximal regions of *SDI1* and *SDI2* promoters

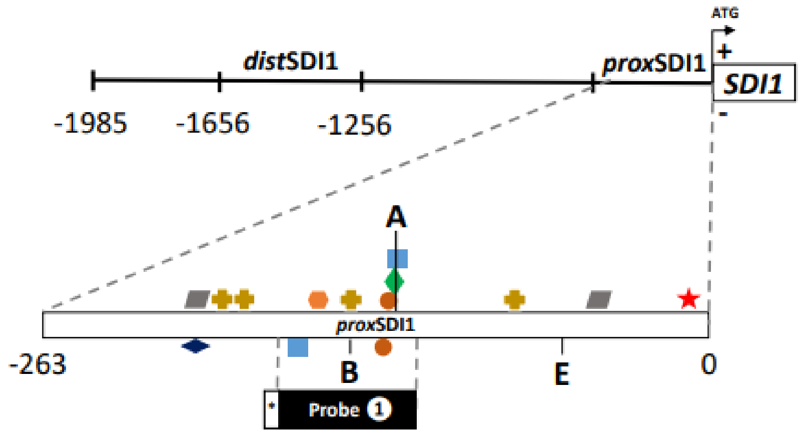


Source for finding a palindromic sequence:
http://bioinformatics.ibp.cz

### Probe 1

| Length – Spacer – Mismatches | Sequence |
|---|---|
| 6 – 1 – 1 | SURE E box bZIP core TEBS TEBS SURE<br>GTGGTAAGAAGTCTCCAAGTGACGTGGCAGCTTCTATGAACAGAGACGAA |
| 6 – 2 – 1 | SURE E box bZIP core TEBS TEBS SURE<br>GTGGTAAGAAGTCTCCAAGTGACGTGGCAGCTTCTATGAACAGAGACGAA |

**Figure S3:** Palindromic sequences within proximal regions of *SDI1* and *SDI2* promoters

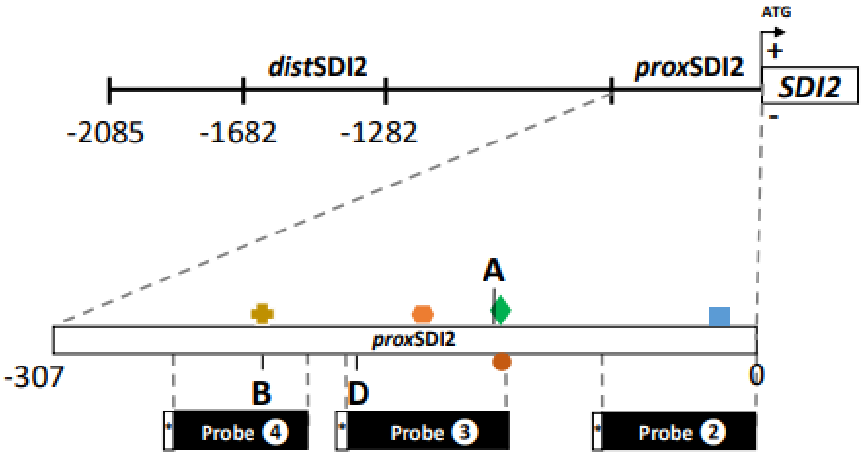


### Probe 2

| Length – Spacer – Mismatches | |
|---|---|
| 8 – 3 – 1 | SURE<br>GAGAG**ATAAAGAA**GGT**TTCTTCAT**TTAGAAACA<u>GAGAC</u>ACGAAGTTTGCG |
| 7 – 4 – 1 | SURE<br>GAGAGAT**AAAGAAG**GTTT**CTTCATT**TAGAAACA<u>GAGAC</u>ACGAAGTTTGCG |
| 6 – 6 – 0 | SURE<br>GAGAGATAAAGAAG**GTTTCT**TCATTT**AGAAAC**A<u>GAGAC</u>ACGAAGTTTGCG |

### Probe 3

No palindromic sequence

### Probe 4

| Length – Spacer – Mismatches | |
|---|---|
| 6 – 7 – 1 | bZIP core<br>GACGG**ATAGGG**T<u>ACGT</u>GG**CACTAT**TTTATA |
