## Supplementary material for "*In silico* analysis of *cis*-elements and identification of transcription factors putatively involved in the regulation of the OAS cluster genes *SDI1* and *SDI2*": Figure S4

**Figure S4:** Western blotting of the candidate TFs fused to the GST protein

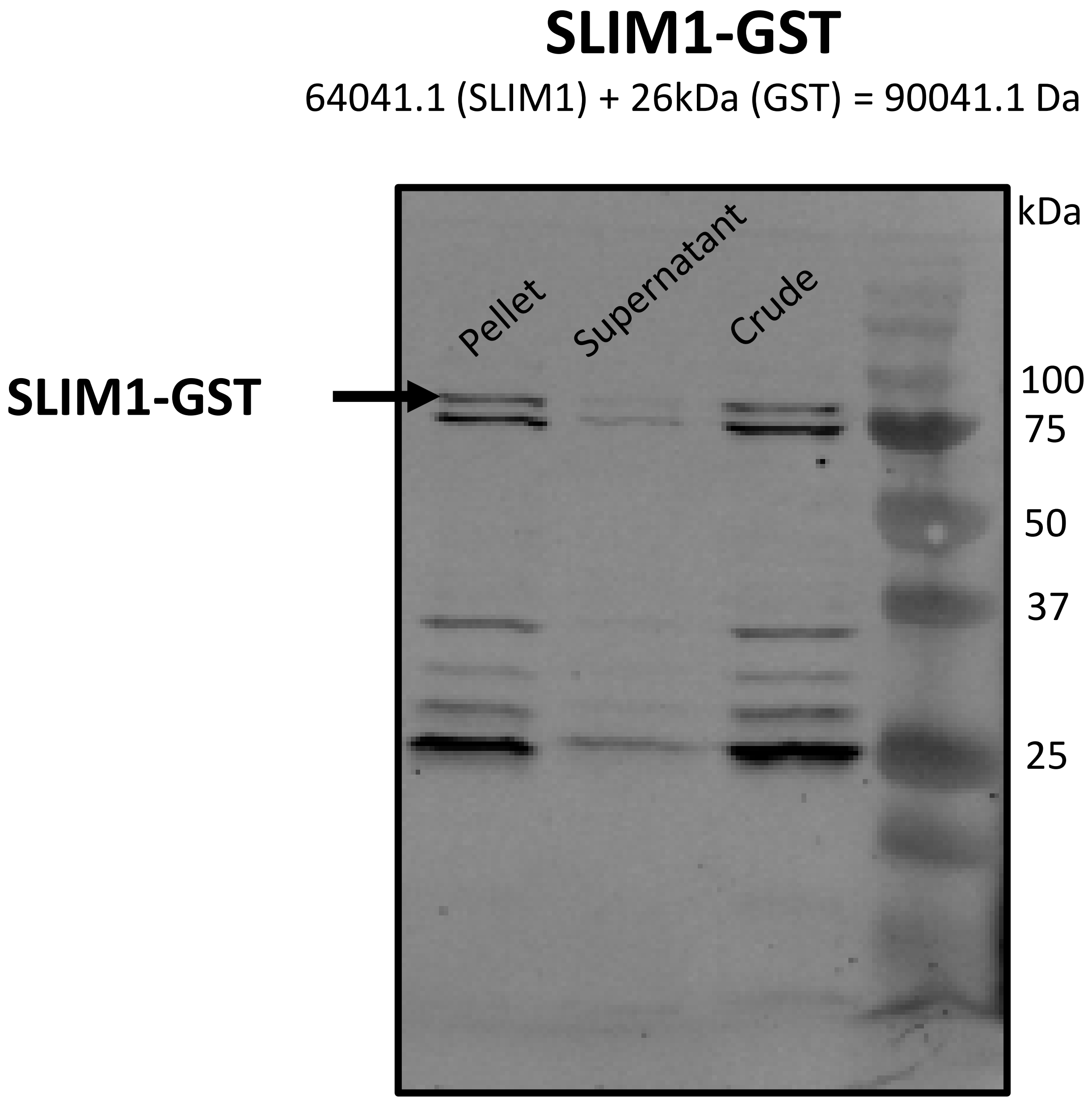

### GBF1-GST

33932.1 (GBF1) + 26kDa (GST) = 59932.1 Da

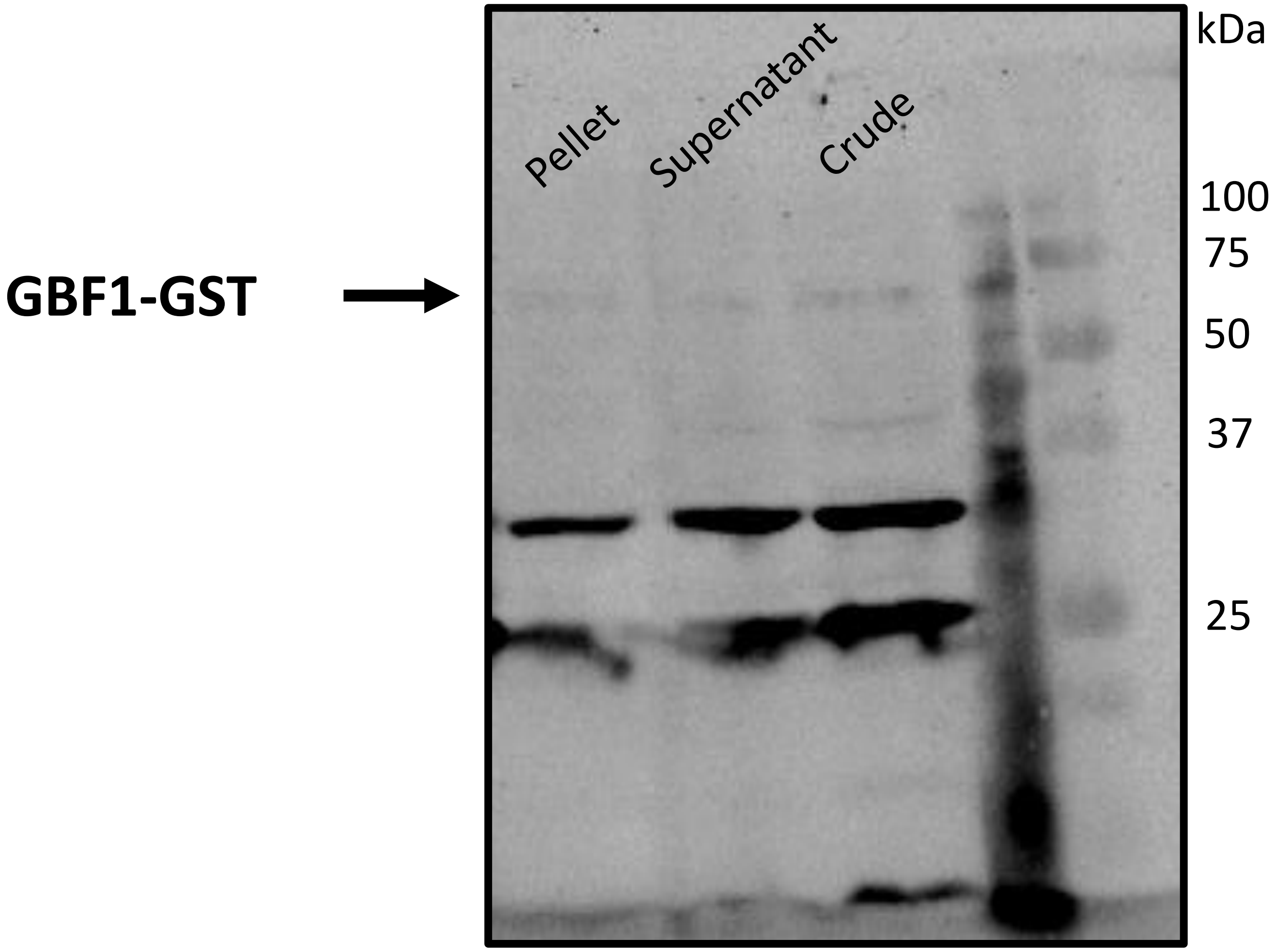

### HYH-GST

16898.5 (HYH)   + 26kDa (GST) = 42898.5 Da

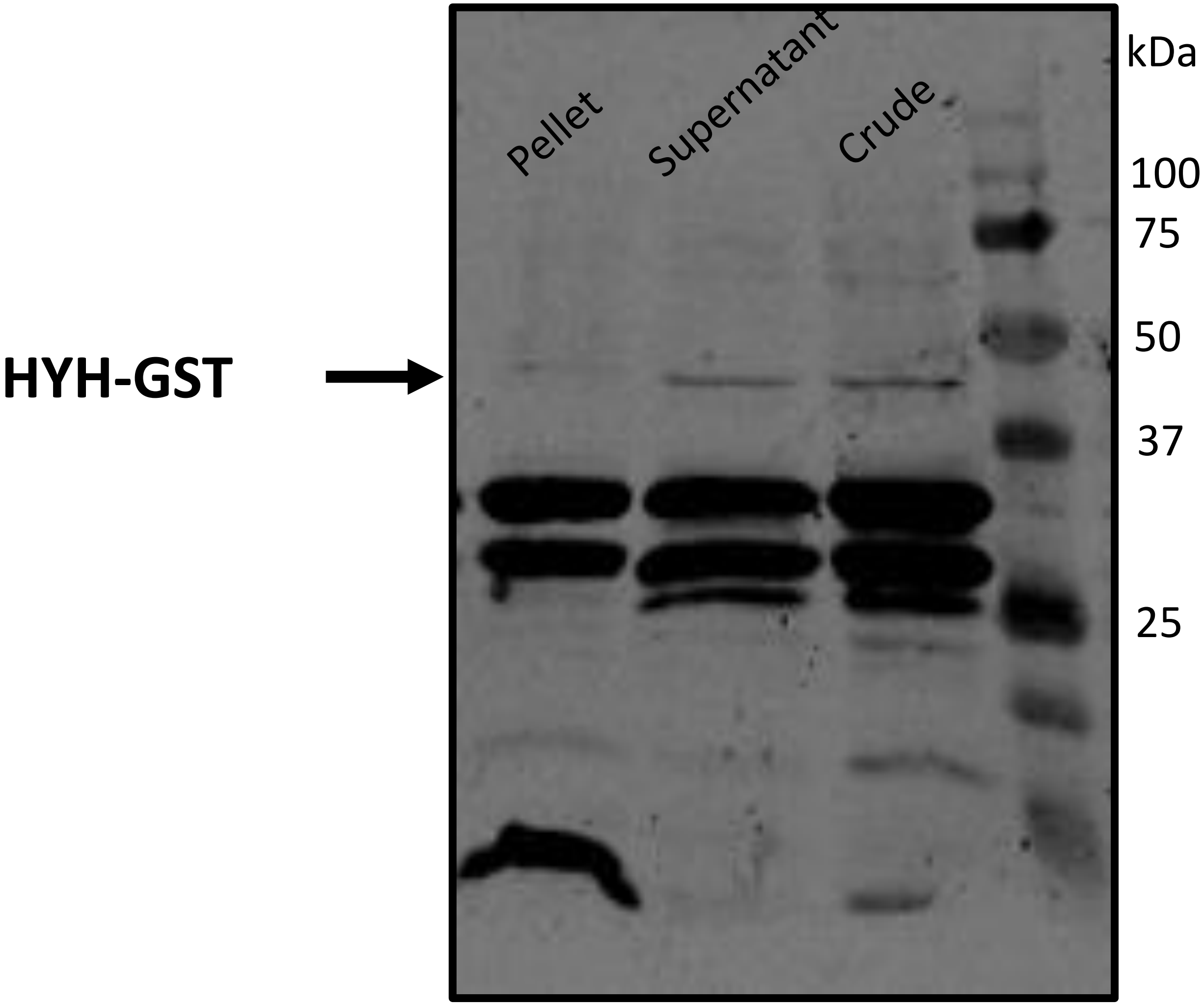

### bZIP16-GST

43109.1 (bZIP16) + 26kDa (GST) = 69109.1 Da

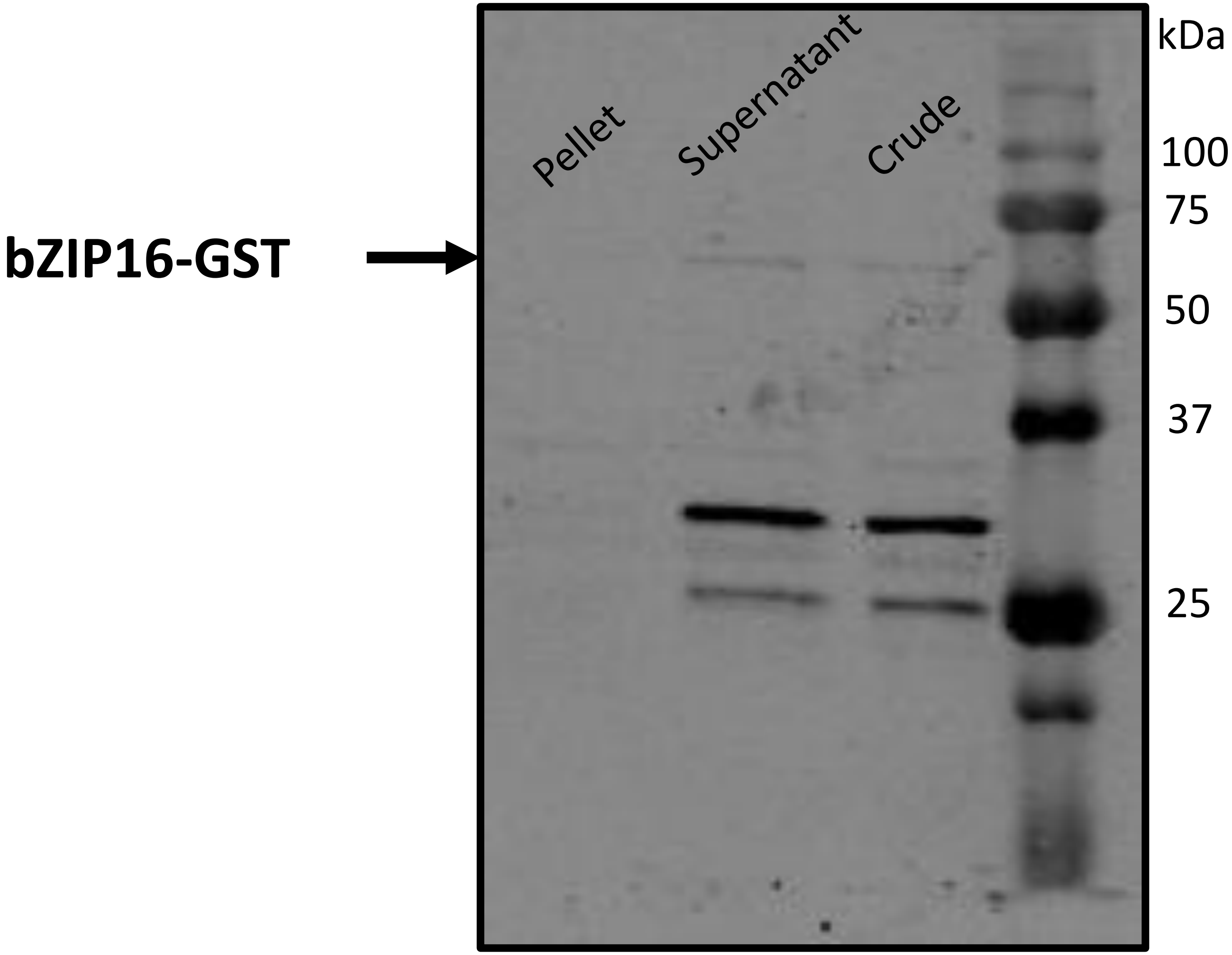

### bZIP44-GST

19073.1 (bZIP44) + 26kDa (GST) = 45073.1 Da

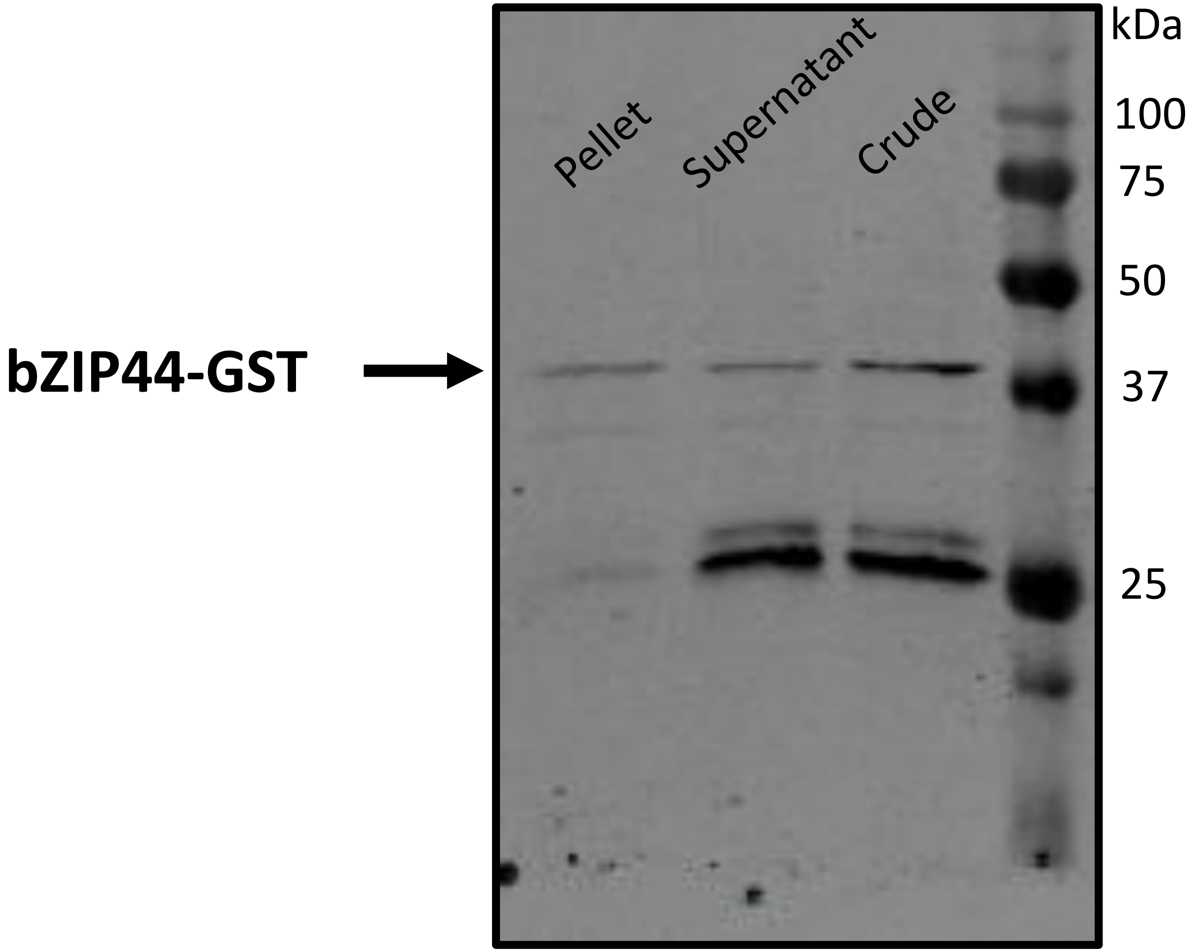

### GST (Entry vector)

26 kDa

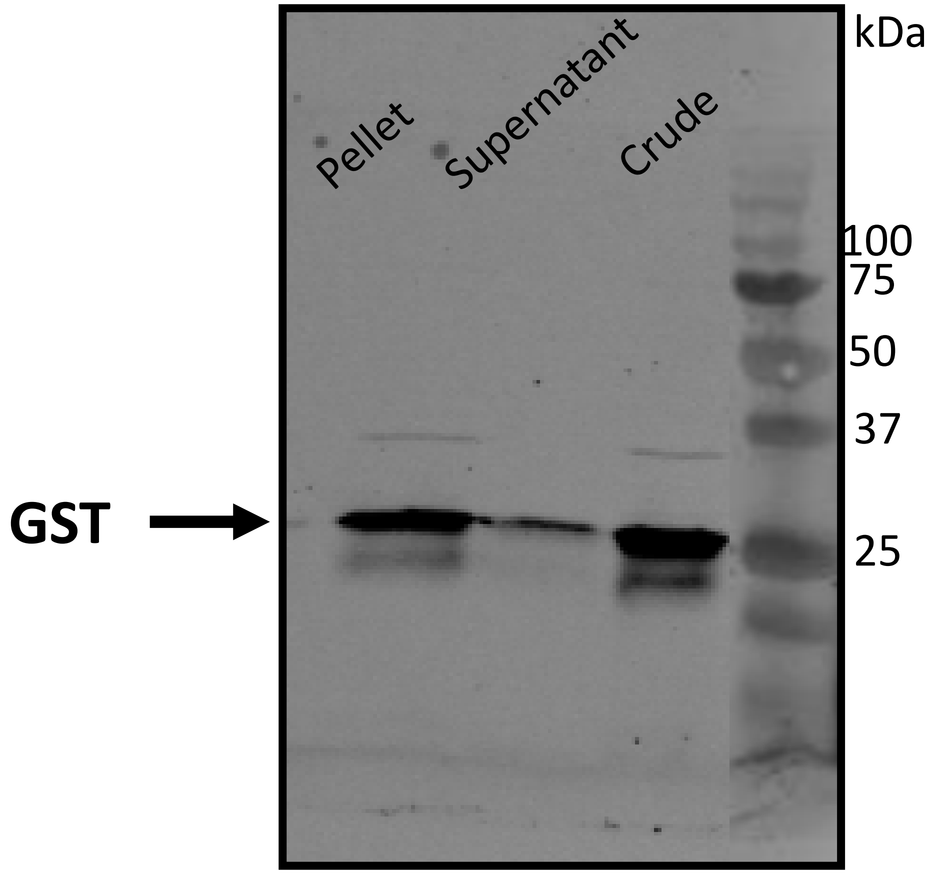
