## Supplementary material for "*In silico* analysis of *cis*-elements and identification of transcription factors putatively involved in the regulation of the OAS cluster genes *SDI1* and *SDI2*": Table S3

**Table S3: List of TFs predicted by FootprintDB to bind to putative motifs A through E**

**Remarks:**

The STAMP E-value threshold in this analysis was 1e-03. This value and the DNA motif similarity score were calculated by BLASTP alignments against the 3D-footprint library (http://floresta.eead.csic.es/3dfootprint/download /list_interface2dna.txt) based on the sum of the Pearson correlation coefficients of the aligned DNA motif positions.

Rows colored white passed the twilight threshold. Rows colored pink did not pass the twilight threshold.

The twilight threshold is associated with identity and similarity thresholds, which were calculated for each sequence alignment by finding cut-off values that increasingly left 95, 90 and 75% of dissimilar TF-DNA complexes below the selected value (for more detail, https://doi.org/10.1093/nar/gks1301; Sebastian and Contreras-Moreira, 2013).

In the footprintDB PWM / Consensus column, the base letters in the upper row indicate the motif’s sequence within the *SDI1* or *SDI2* promoter, whereas the base letters in the lower row indicate the binding site of the TF candidate, which is predicted by footprintDB to potentially bind the motif. Bases that match between the motif and the TF binding site are shown in uppercase, whereas unmatched bases are shown in lowercase.

The base letters presented follow the IUPAC nucleotide code:

| IUPAC nucleotide code | Base |
| --- | --- |
| A | Adenine |
| C | Cytosine |
| G | Guanine |
| T | Thymine |
| R | A or G |
| Y | C or T |
| S | G or C |
| W | A or T |
| K | G or T |
| M | A or C |
| B | C or G or T |
| D | A or G or T |
| H | A or C or T |
| V | A or C or G |
| N | any base |
| - | gap |

**footprintDB results for Motif A**

Query:  [TGAACAGAGACG](http://floresta.eead.csic.es/footprintdb/results/765711c6f5b5eeb929452622a4e164b1/Motif%20A.data.transfac.html#DNA) (SDI1)

Query:  [TGAACCAAGACG](http://floresta.eead.csic.es/footprintdb/results/2e7b8701687ec800da34effa74daf78c/Motif%20A_SDI2.data.transfac.html#DNA) (SDI2)

| ***In silico* TF candidate** | **Source** | **STAMP e-value** | **Motif similarity** | **footprinDB PWM / Consensus** |
| --- | --- | --- | --- | --- |
| EIN3 | AthalianaCistrome v4_May2016 | 3.8e-07 | 8.36 / 11 | ----TGAACCAAGACG [wCAwTGwAyyTrgaC-](http://floresta.eead.csic.es/footprintdb/index.php?motif=28481) |
| REF6 | JASPAR 2020 | 2.3e-05 | 7.00 / 11 | CGTCTCTGTTCA [-trCTCTGTTTy](http://floresta.eead.csic.es/footprintdb/index.php?motif=25421) |
| SLIM1 | AthalianaCistrome v4_May2016 | 4.1e-05 | 7.86 / 12 | -CGTCTCTGTTCA-------- [ryGTCyAGrTtCAwwrwatht](http://floresta.eead.csic.es/footprintdb/index.php?motif=28482) |
| GAL1 | AthalianaCistrome v4_May2016 | 5.2e-05 | 6.84 / 10 | TGAACCAAGACG- [--rwCGrygaCGw](http://floresta.eead.csic.es/footprintdb/index.php?motif=28442) |
| ANAC2 | UniPROBE 20160601 | 7.4e-05 | 6.98 / 8 | ----CGTCTCTGTTCA [mrCACGtCTCTy----](http://floresta.eead.csic.es/footprintdb/index.php?motif=17386) |
| GAL3 | AthalianaCistrome v4_May2016 | 1.0e-04 | 7.32 / 12 | --TGAACCAAGACG- [ygaarwCGAwgwCGw](http://floresta.eead.csic.es/footprintdb/index.php?motif=28438) |
| GT-1 | JASPAR 2020 | 3.3e-04 | 5.64 / 7 | -TGAACCAAGACG [gTTAACCa-----](http://floresta.eead.csic.es/footprintdb/index.php?motif=13492) |
| ARR11 | JASPAR 2020 | 4.2e-04 | 5.59 / 8 | CGTCTCTGTTCA [CGTATCTw----](http://floresta.eead.csic.es/footprintdb/index.php?motif=25070) |
| HAT3.1 | CISBP 1.02 | 4.8e-04 | 6.14 / 9 | -TGAACCAAGACG [sTrmACcahy---](http://floresta.eead.csic.es/footprintdb/index.php?motif=11345) |
| NAC68 | JASPAR 2020 | 0.001 | 7.21 / 12 | ---------TGAACCAAGACG--- [wwtaACTTGTgwwmyAAGTAAcww](http://floresta.eead.csic.es/footprintdb/index.php?motif=33271) |

**footprintDB results for Motif B**

Query:  [GTGACGTGGCA](http://floresta.eead.csic.es/footprintdb/results/02ae0aa9668179871c150aa3fcf98ccf/Motif%20B_SDI1.data.transfac.html#DNA) (SDI1)

Query:  [GGTACGTGGCA](http://floresta.eead.csic.es/footprintdb/results/d035726526439601f71d60c7bb15b017/Motif%20B_SDI2.data.transfac.html#DNA) (SDI2)

| ***In silico* TF candidate** | **Source** | **STAMP e-value** | **Motif similarity** | **footprinDB PWM / Consensus** |
| --- | --- | --- | --- | --- |
| bZIP60 | ArabidopsisPBM 20140210 | 5.6e-16 | 9.58 / 10 | GTGACGTGGCA [rTGACGTGgc-](http://floresta.eead.csic.es/footprintdb/index.php?motif=7290) |
| ABF4 | JASPAR 2020 | 2.6e-15 | 9.96 / 11 | -GTGACGTGGCA [drwsACGTGGma](http://floresta.eead.csic.es/footprintdb/index.php?motif=32960) |
| HYH | JASPAR 2020 | 7.7e-15 | 9.85 / 11 | -GTGACGTGGCA [grwsACGTGkca](http://floresta.eead.csic.es/footprintdb/index.php?motif=32853) |
| ABI5 | AthalianaCistrome v4_May2016 | 1.0e-14 | 10.21 / 11 | ----GTGACGTGGCA rrwGrwsACGTGGCA |
| bZIP42 | AthalianaCistrome v4_May2016 | 1.4e-14 | 9.94 / 11 | --GTGACGTGGCA [TGCTGACGTGGCa](http://floresta.eead.csic.es/footprintdb/index.php?motif=28381) |
| bZIP16 | [JASPAR 2020](http://jaspar.genereg.net/) AthalianaCistrome v4_May2016 | 3.8e-14 | 10.07 / 11 | ----GTGACGTGGCA [dwwksysACGTGGCA](http://floresta.eead.csic.es/footprintdb/index.php?motif=25355) |
| GBF2 | JASPAR 2020 | 4.2e-14 | 9.82 / 11 | GTGACGTGGCA-- [rhsACGTGGCAww](http://floresta.eead.csic.es/footprintdb/index.php?motif=32968) |
| ABI5 | AthalianaCistrome v4_May2016 | 1.1e-13 | 10.16 / 11 | -----GTGACGTGGCA-- [wdrwgrwsACGTGKCarw](http://floresta.eead.csic.es/footprintdb/index.php?motif=28364) |
| GBF3 | [JASPAR 2020](http://jaspar.genereg.net/) AthalianaCistrome v4_May2016 | 1.2e-13 | 9.95 / 11 | ----GTGACGTGGCA [dwwkstsACGTGGCA](http://floresta.eead.csic.es/footprintdb/index.php?motif=25357) |
| TOE1 | ArabidopsisPBM 20140210 | 4.9e-08 | 7.54 / 10 | GGTACGTGGCA [gGTACGAGGW-](http://floresta.eead.csic.es/footprintdb/index.php?motif=7271) |
| GBF1 | CISBP 1.02 | 5.5e-08 | 7.52 / 10 | GGTACGTGGCA [-ysACGTGkmm](http://floresta.eead.csic.es/footprintdb/index.php?motif=10806) |
| PIF4 | JASPAR 2020 | 1.2e-07 | 6.78 / 8 | GGTACGTGGCA [--CACGTGsc-](http://floresta.eead.csic.es/footprintdb/index.php?motif=9044) |
| ABF1 | JASPAR 2020 | 1.3e-07 | 7.85 / 10 | GGTACGTGGCA-- [-aCACGTGkCAww](http://floresta.eead.csic.es/footprintdb/index.php?motif=32783) |

**footprintDB results for Motif C**

Query:   [ACCTACGTGGTA](http://floresta.eead.csic.es/footprintdb/results/471666969a67698fa12cbd319e65a067/Motif%20C_SDI1.data.transfac.html#DNA) (SDI1)

Query:   [TAACACGTAGGC](http://floresta.eead.csic.es/footprintdb/results/39e696864f85961932bad0c873f98b65/Motif%20C_SDI2.data.transfac.html#DNA) (SDI2)

| ***In silico* TF candidate** | **Source** | **STAMP e-value** | **Motif similarity** | **footprinDB PWM / Consensus** |
| --- | --- | --- | --- | --- |
| NAC055 | JASPAR 2020 | 6.8e-08 | 6.95 / 8 | TAACACGTAGGC [----ACACGTAA--](http://floresta.eead.csic.es/footprintdb/index.php?motif=25068) |
| PIF3 | ArabidopsisPBM 20140210 | 1.3e-06 | 7.22 / 10 | ACCTACGTGGTA [-sCCACGTGGs-](http://floresta.eead.csic.es/footprintdb/index.php?motif=7247) |
| TOE2 | ArabidopsisPBM 20140210 | 3.0e-06 | 7.08 / 8 | --ACCTACGTGGTA [wAACCTACGw----](http://floresta.eead.csic.es/footprintdb/index.php?motif=7275) |
| MYC2 | 3D-footprint 20200120 | 5.2e-06 | 5.97 / 7 | TAACACGTAGGC [--aCaCGtG---](http://floresta.eead.csic.es/footprintdb/index.php?motif=32438) |
| SPT | CISBP 1.02 | 9.3e-06 | 6.60 / 9 | ACCTACGTGGTA [-csCACGTGc--](http://floresta.eead.csic.es/footprintdb/index.php?motif=10713) |
| ABF1 | CISBP 1.02 | 1.2e-05 | 7.11 / 11 | TAACACGTAGGC [graCACGTGkb-](http://floresta.eead.csic.es/footprintdb/index.php?motif=10796) |
| TOE1 | ArabidopsisPBM 20140210 | 1.6e-05 | 6.80 / 8 | --ACCTACGTGGTA [rAACcTrsGA----](http://floresta.eead.csic.es/footprintdb/index.php?motif=7272) |
| HY5 | Athamap 20091028 | 3.6e-05 | 6.91 / 10 | ACCTACGTGGTA- [--CCACGTGkCAT](http://floresta.eead.csic.es/footprintdb/index.php?motif=5026) |
| ABI5 | Athamap 20091028 | 4.4e-05 | 6.34 / 9 | ACCTACGTGGTA [-KAyACGTbR--](http://floresta.eead.csic.es/footprintdb/index.php?motif=4997) |

**footprintDB results for Motif D**

Query:   [ACCGCGGGCTGG](http://floresta.eead.csic.es/footprintdb/results/ed2b08c50e89f57c152871e74d6265f6/Motif%20D.data.transfac.html#DNA) (SDI2)

| ***In silico* TF candidate** | **Source** | **STAMP e-value** | **Motif similarity** | **footprinDB PWM / Consensus** |
| --- | --- | --- | --- | --- |
| ERF105 | JASPAR 2020 | 3.1e-04 | 5.65 / 8 | ACCGCGGGCTGG [--CGCCGGCm--](http://floresta.eead.csic.es/footprintdb/index.php?motif=13473) |
| ERF15 | CISBP 1.02 | 7.6e-04 | 5.48 / 8 | ACCGCGGGCTGG [--cGsCGGCc--](http://floresta.eead.csic.es/footprintdb/index.php?motif=10580) |
| ERF6 | JASPAR 2020 | 0.001 | 5.96 / 10 | ACCGCGGGCTGG [-vcGCCGGCak-](http://floresta.eead.csic.es/footprintdb/index.php?motif=13478) |
| ERF4 | JASPAR 2020 | 0.002 | 5.33 / 8 | ACCGCGGGCTGG [-cCGCCGcC---](http://floresta.eead.csic.es/footprintdb/index.php?motif=13465) |
| TOE2 | ArabidopsisPBM 20140210 | 0.002 | 5.87 / 10 | ACCGCGGGCTGG [wCCTCGTACt--](http://floresta.eead.csic.es/footprintdb/index.php?motif=7274) |

**footprintDB results for Motif E**

Query: [GGGAGAGACAAC](http://floresta.eead.csic.es/footprintdb/results/c2db0457cb9f81c788578e38fcad74ac/Motif%20E.data.transfac.html#DNA) (SDI1)

| ***In silico* TF candidate** | **Source** | **STAMP e-value** | **Motif similarity** | **footprinDB PWM / Consensus** |
| --- | --- | --- | --- | --- |
| ARF1 | 3D-footprint 20200120 | 6.3e-06 | 6.67 / 9 | GGGAGAGACAAC [--GGGAnACAA-](http://floresta.eead.csic.es/footprintdb/index.php?motif=32217) |
| BPC1 | AthalianaCistrome v4_May2016 | 6.2e-05 | 7.43 / 12 | GGGAGAGACAAC--- [rAGAGAGAGArAGAg](http://floresta.eead.csic.es/footprintdb/index.php?motif=28329) |
| FRS9 | AthalianaCistrome v4_May2016 | 2.6e-04 | 7.48 / 12 | --GGGAGAGACAAC------- [GAGAGAGAGAGAGAGAGAGAG](http://floresta.eead.csic.es/footprintdb/index.php?motif=28744) |
| ANAC2 | UniPROBE 20160601 | 4.6e-04 | 6.59 / 11 | GGGAGAGACAAC- [-raAGAGmCGTAt](http://floresta.eead.csic.es/footprintdb/index.php?motif=17385) |
| WRKY23 | JASPAR 2020 | 5.2e-04 | 5.55 / 7 | GGGAGAGACAAC- [-----RGTCAACG](http://floresta.eead.csic.es/footprintdb/index.php?motif=13550) |
| BPC6 | [JASPAR 2020](http://jaspar.genereg.net/) AthalianaCistrome v4_May2016 | 5.3e-04 | 7.32 / 12 | ----GGGAGAGACAAC----- [kAGAGAGAGAGAGAGAGAGAG](http://floresta.eead.csic.es/footprintdb/index.php?motif=25408) |
| WRKY30 | AthalianaCistrome v4_May2016 | 6.2e-04 | 6.32 / 9 | GGGAGAGACAAC-- [---aAAGTCAACGc](http://floresta.eead.csic.es/footprintdb/index.php?motif=28825) |
