## Supplementary material for "*In silico* analysis of *cis*-elements and identification of transcription factors putatively involved in the regulation of the OAS cluster genes *SDI1* and *SDI2*": Table S6

**Table S6**: **Sequence context of the most proximal SURE element in the promoters of two presumed SLIM1-target genes lacking a UPE box, UPE-like motif or TEBS in their promoters**

| **AGI** | **Promoter sequence context of the most proximal SURE element** | **Location of SURE relative to ATG** |
| --- | --- | --- |
| AT2G44460 | **ACGTAG**ACCAA**CAACGT**CACGAGACTAAGCTC | -424 |
| AT5G54390 | GTCTC**TCATCA**TCCG**TGTTGA** | -1349 |

**Remarks**: The intersection between SLIM1-dependent lowS-responsive genes (Maruyama-Nakashita et al., 2006) and genes with SLIM1 binding peaks by DAP-seq (O’Malley et al., 2016) contains 14 genes. The promoters of all 14 of those genes contain at least 1 SURE element. Two of the 14 genes do not contain a UPE-like motif, a UPE box, or a TEBS element. The sequence context around the most proximal SURE element in the promoters of these two genes is shown above. Palindromic sequence is in bold text; SURE elements are underlined.
